# Dietary serine protects the retinal pigmented epithelium by blunting reactive oxygen species in dry age-related macular degeneration

**DOI:** 10.64898/2026.04.30.722030

**Authors:** Ganesh Satyanarayana, Rabina Kumpakha, Jack Papania, Jana Sellers, Micah Chrenek, James T. Handa, Sayantan Datta

## Abstract

Age-related macular degeneration (AMD) is a progressive complex eye disease and one of the leading causes of blindness. AMD progression is marked by molecular changes in the retinal pigmented epithelium (RPE) which include increased reactive oxygen species (ROS) accumulation, mitochondrial dysfunction - eventually leading to dysfunctional RPE. Mitophagy regulator, Pink1, is reduced in the RPE of AMD patients and Pink1 loss leads to a shift from mitochondrial respiration to glycolysis. Serine is a non-essential amino acid which is *de novo* synthesized from glycolytic intermediate 3-PG via the rate limiting enzyme PHGDH. Serine is tightly integrated into anabolic processes like glutathione (GSH) cycling, maintaining NADH/NADPH pools leading to changes in AMPK signaling. Here, we show that Pink1 loss leads to a reduction in PHGDH and serine levels in the RPE leading to impaired mitochondrial structure and function, increased ROS mediated damage, increased inflammation, and hampered retinal function. Serine supplementation rescued ROS accumulation, balanced GSH abundance, and increased retinal function. Overall, our study highlights the potential of dietary serine in ROS management in AMD.

## Introduction

Age-related macular degeneration (AMD) remains the leading cause of irreversible blindness in the elderly population^1^. AMD pathology manifests primarily in the macula, where the degeneration of the retinal pigment epithelium (RPE) and subsequent loss of photoreceptors characterize the major non-exudative (dry) form of the disease and has limited treatment options.^2^. RPE dysfunction is an early feature of AMD showing hallmark characteristics of reactive oxygen species (ROS) accumulation mediated oxidative stress leading to mitochondrial dysfunction, failure of cytoprotective mechanisms and oxidative stress. Due to its high metabolic activity, the RPE is uniquely vulnerable to reactive oxygen species (ROS) accumulation^3,4^. To maintain homeostasis under such a high oxidative load, the RPE relies heavily on mitochondrial quality control called mitophagy. Mitochondrial PINK1 (PTEN-induced kinase 1) acts as a master regulator of mitophagy and helps maintain mitochondrial homeostasis by aiding in the clearance of dysfunctional mitochondria. Under Pink1 loss, this clearance mechanism fails, damaged mitochondria accumulate thereby perpetuating a cycle of ROS accumulation and ATP depletion that drives RPE dysfunction ^6,7^.

In recent studies, among mitochondrial quality control proteins, PINK1 exhibited the most significant reduction in the primary retinal pigment epithelium (RPE) from donor eyes of AMD patients^8^. This loss of PINK1 compromises mitophagy, leading to a metabolic shift towards glycolysis, ROS accumulation and eventual problems with RPE structure and function ^3,9^.

Serine synthesis pathway (SSP) is an offshoot of glycolysis with PHGDH driving glycolytic flux from 3-phosphoglycerate to 3-phosphopyruvate. PHGDH is the rate limiting step in the serine synthesis pathway (SSP) from glycolysis^10,11^. Serine is considered a non-essential amino acid and has a canonical role in anabolic processes. However, recent studies have shown exogenous serine supplementation can aid in supporting antioxidant intermediates such as glutathione (GSH) and NADPH and NADH^12-14^. Furthermore, other studies have also shown the potential of serine to activate key cytoprotective pathways via AMPK and mTOR signaling^15-16^. Serine insufficiency is one of the drivers for the advancement of macular telangiectasia type 2 (MACTEL) wherein non-functional PHGDH mutations reduce serine levels in the retinas of these patients^16.^ Serine supplementation is well tolerated and safe up to 400mg/Kg/day making it an easily translatable treatment option for early AMD. Beyond retinal diseases, serine supplementation is also being tested as a possible therapeutic in Alzheimer’s disease and hereditary sensory autonomic neuropathy type I (HSAN1) where serine deficiency has not been implicated (NCT01733407; NCT06113055). Increased serine metabolism provides antioxidant support by aiding in the production of metabolites glutathione (GSH), nicotinamide adenine dinucleotide phosphate (NADPH) which facilitate rapid ROS quenching in both cytoplasmic and mitochondrial compartments. These studies lend credence to the notion that serine supplementation can have a wide range of benefits and cytoprotective effects pointing towards it being a potential supplement to protect the RPE during AMD pathogenesis.

To define how loss of PINK1 disrupts retinal pigment epithelium (RPE) homeostasis, we examined the relationship between mitochondrial dysfunction, serine biosynthesis, and redox control under PINK1-deficient conditions. Given the central role of Pink1 in maintaining mitochondrial integrity and quality control, we hypothesized that Pink1 loss would destabilize mitochondrial homeostasis. Interestingly, loss of Pink1 led to impaired serine synthesis pathway, a key contributor to glutathione mediated antioxidant capacity. We assessed alterations in mitochondrial function, oxidative stress, and inflammatory signaling followed by retinal functional tests *in vivo*. We further tested whether exogenous serine supplementation could rescue mitochondrial homeostasis, ROS accumulation, and retinal function. Together, these studies establish Pink1 as a critical upstream regulator of serine metabolism and demonstrate that dietary serine supplementation provides robust support against ROS imbalance in the aging RPE.

In this study, we explore the mechanistic consequences of PINK1 loss in the RPE and utilize serine as a dietary supplement for quenching ROS accumulation and rescuing RPE and retinal structure and function.

## Results

### PINK1 and PHGDH are reduced in AMD Patient iPSC-RPE

We analyzed PHGDH in retinal tissue sections from an 86-year-old AMD patient donor. Immunofluorescence staining of paraffin-embedded human sections revealed a marked reduction in PHGDH expression in the RPE, with signal intensity reduced by ∼60% compared to adjacent, unaffected regions (**Fig 1A**). Next, we tested iPSC-RPE derived from AMD patients classified as dry and advanced AMD for PHGDH mRNA and protein levels. PHGDH expression was reduced in patient iPSC-RPE cells (**Fig 1B, C**) when compared to age-matched controls. Interestingly, the secondary genes in the SSP remained unchanged in these cells by ∼50% compared to non-diseased control iPSC-RPE (**Fig S1A-D**). We have previously reported that Pink1 was reduced in the RPE cells over drusen. Consistently, we detected a significant reduction of ∼50% in PINK1 mRNA levels in AMD patient iPSC-RPE cells compared to non-diseased controls indicating that mitochondrial quality control defects are possibly intrinsic in the aging RPE (**Fig 1D, E**). Overall, Pink1 loss potentially contributes to loss of PHGDH in AMD patient tissue sections and patient derived iPSC-RPE cells.

**Figure 1.**
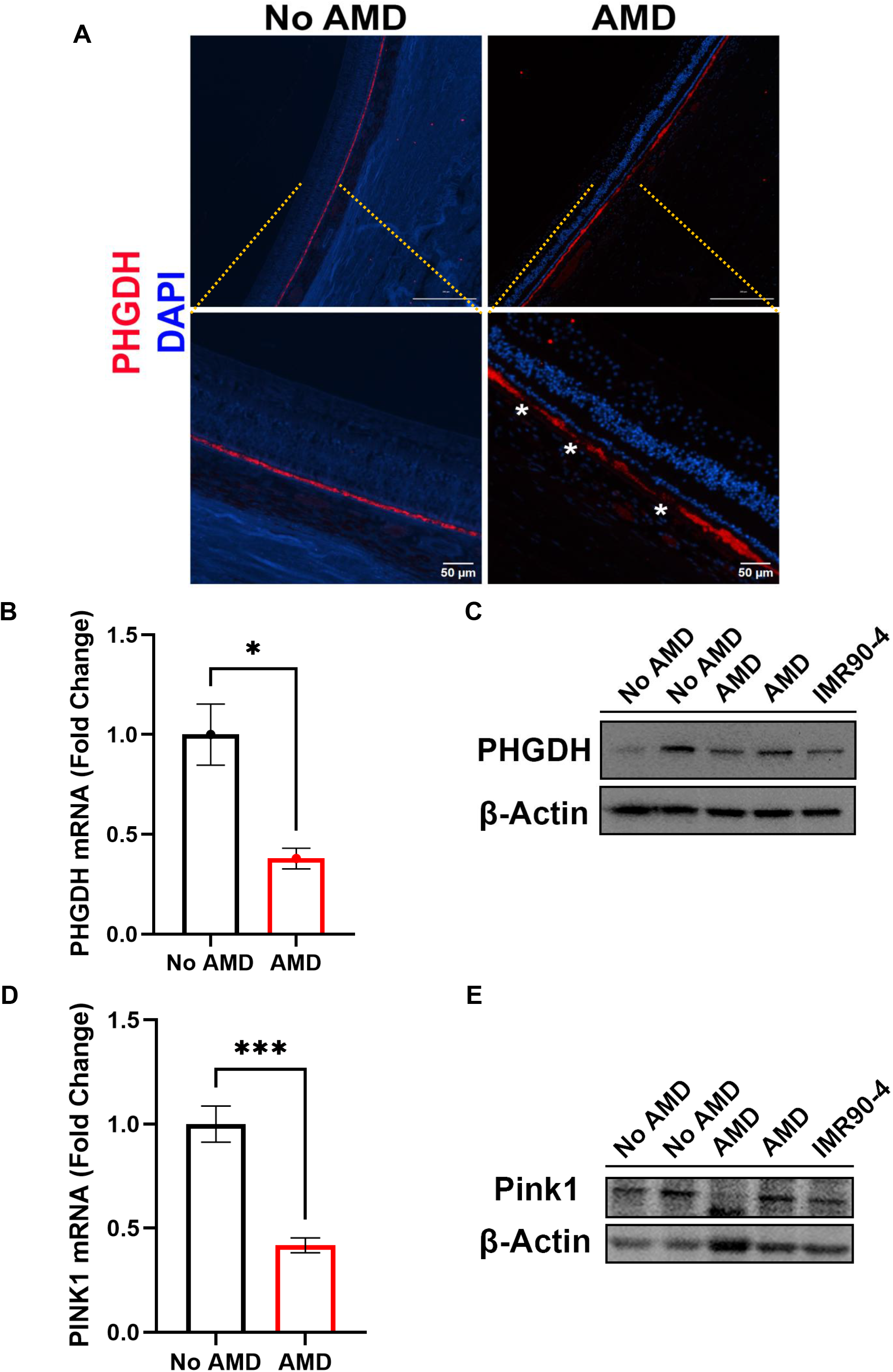
AMD patient tissues and iPSC-RPE have reduced PHGDH and PINK1. (**A**) Retinal sections from an AMD patient aged 87yo and non-diseased control eyes were processed and stained with PHGDH (CST13428) immunofluorescence. White stars indicate areas under drusen under the peripheral RPE. (**B**) 10ng of cDNA was loaded for qPCR analyses using SYBR Green kits (NEB M3003) and (**C**) 10ug of cell lysates were loaded for western blots for PHGDH (CST13428). PINK1 levels were assessed with (**D**) qPCR and in (**E**) western blots (SC33796). qPCR data are presented as 6 technical replicates for each patient. Statistical significance was determined using two-way ANOVA with siPINK1 (knockdown vs. control) and +/- serine supplementation, followed by Tukey’s post hoc multiple comparisons test. (*p<0.05; ***p<0.0005;)1

### PINK1 loss disrupts serine biosynthesis

To determine whether PINK1 loss directly reduces PHGDH expression and serine levels in the RPE, ARPE-19 cells were treated with siPINK1 for 5 days with or without L-serine supplementation (200 µM; **Fig. 2A**). PHGDH mRNA and protein levels were significantly reduced by ∼85% in siPINK1 cells and were not restored by serine supplementation (**Fig. 2B–D**). Expression of other serine synthesis pathway genes (PSAT1, PSPH, SHMT1, and SHMT2) was not significantly affected by PINK1 loss (**Fig. S2A–E**). Intracellular serine levels increased by 1.5-fold via exogenous supplementation, confirming effective cellular uptake. Seahorse extracellular flux analysis revealed that basal oxygen consumption rate and maximal respiratory capacity were reduced by ∼25% in siPINK1 cells and were not rescued by serine supplementation (**Fig. 2E–G**). Cells deficient in PINK1 exhibited a shift toward glycolytic energy metabolism (**Fig. 2H**), while serine supplementation in PINK1-sufficient cells increased reliance on oxidative phosphorylation. Cellular ATP levels were reduced by ∼20% in siPink1 cells and were restored back to untreated levels by serine supplementation (**Fig. S2F**). Together, these data indicate that PINK1 deficiency impairs both PHGDH-dependent serine biosynthesis and mitochondrial respiration through mechanisms that are not alleviated by exogenous serine.

**Figure 2.**
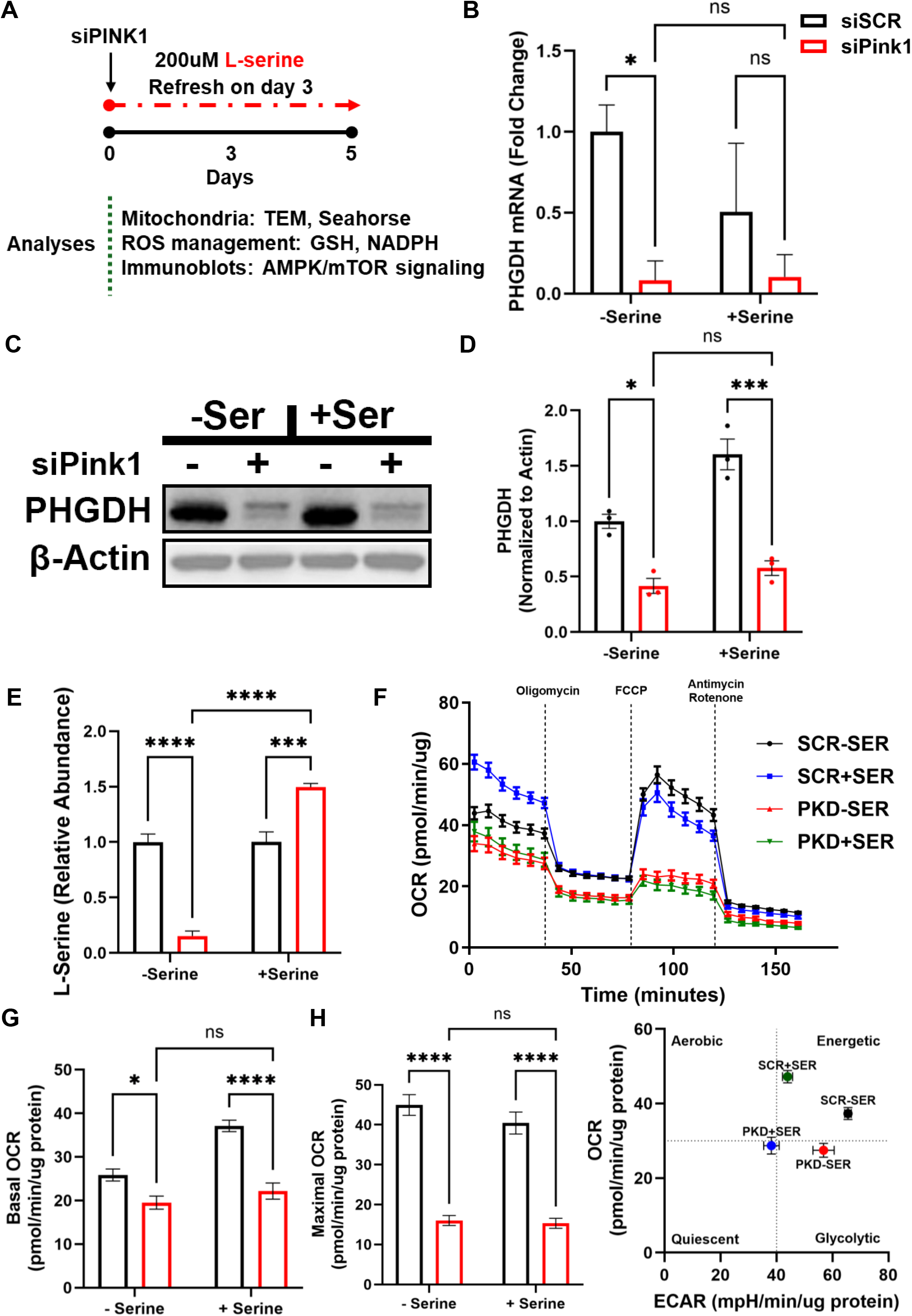
PINK1 loss reduces PHGDH and blunts mitochondrial respiration. **(A)** ARPE-19 cells were treated with 20nmols of siScramble or siPink1 for 5 days. Cells were supplemented with 200uM serine refreshed at 3 days. **(B)** 10ng of cDNA was loaded for qPCR analyses using SYBR Green kit (NEB M3003) for PHGDH. **(C, D)** 10ug of cell lysates were loaded for western blots for PHGDH (CST13428) and quantified. **(E)** L-serine levels were measured using a kit (Sigma MAK352). **(F)** ARPE-19 cells were treated with either siPINK1 or siScramble as mentioned. 40,000 cells were plated on Seahorse XF Pro plates. MitoStress test was performed to quantify oxygen consumption rate for siPINK1 ± serine supplementation. **(G)** Basal OCR was assessed within the first 40 minutes of the test. **(H)** Maximal respiratory potential was analyzed after FCCP treatment at 60 minutes. **(I)** Cellular energy profile was assessed by plotting OCR and ECAR to identify differential glycolysis and mitochondrial respiration in Pink1 deficient ± serine supplemented cells. Data are presented as mean +/- SEM from n = 3 replicates per group for qPCR and western blots and n=6 for Seahorse analyses. Statistical significance was determined using two-way ANOVA with siPINK1 (knockdown vs. control) and +/- serine supplementation, followed by Tukey’s post hoc multiple comparisons test. (*p<0.05; ***p<0.0005; ****p<0.0001)

### Serine supplementation restores redox homeostasis disrupted by PINK1 deficiency

To determine whether serine supplementation reduces oxidative stress resulting from PINK1 loss, mitochondrial superoxide was assessed by MitoSOX staining in siPINK1 ARPE-19 cells. MitoSOX signal was elevated by ∼40% in siPINK1 cells and was reduced untreated by serine supplementation in a dose-dependent manner, with 200 µM for 5 days restoring levels to near-control values (**Fig. 3A, C**; **Fig. S3A**). Mitochondrial membrane potential, assessed by TMRM staining, was significantly reduced by ∼40% in siPINK1 cells and was restored by serine supplementation (**Fig. 3B, D**). Total cellular ROS was significantly elevated by ∼66% in siPINK1 cells and significantly reduced back to untreated levels by 200 µM serine supplementation for 5 days (**Fig. 3E**). Intracellular GSH was decreased by ∼20% siPINK1 cells and was significantly restored by ∼15% serine supplementation; the GSH/GSSG ratio was likewise significantly increased (**Fig. 3F, G**). NADPH levels, which were reduced by ∼17% in siPINK1 cells and were restored by serine supplementation (**Fig. 3H**). To determine whether loss of endogenous serine biosynthesis alone is sufficient to recapitulate these redox defects, ARPE-19 cells were transfected with siPHGDH for 5 days. SHMT2 was upregulated by ∼2-fold in siPHGDH cells (**Fig. S3B**). Total NADH was reduced by ∼33% in siPHGDH cells (**Fig. S3C**), and the mitochondrial and cytoplasmic NAD^+^/NADH ratio was increased (**Fig. S3D**). NADPH levels were reduced by ∼40% in both compartments and were rescued by serine supplementation (**Fig. S3E**). Total GSH was reduced by ∼25%, primarily in the cytoplasmic fraction, while GSSG was modestly elevated, resulting in a significantly reduced GSH/GSSG ratio (**Fig. S3F–H**). These data indicate that serine supplementation rescues redox mediators and balances ROS homeostasis brought about by PINK1 loss. These data indicate that PHGDH loss alone is sufficient to impair glutathione homeostasis and NADPH production in RPE cells.

**Figure 3.**
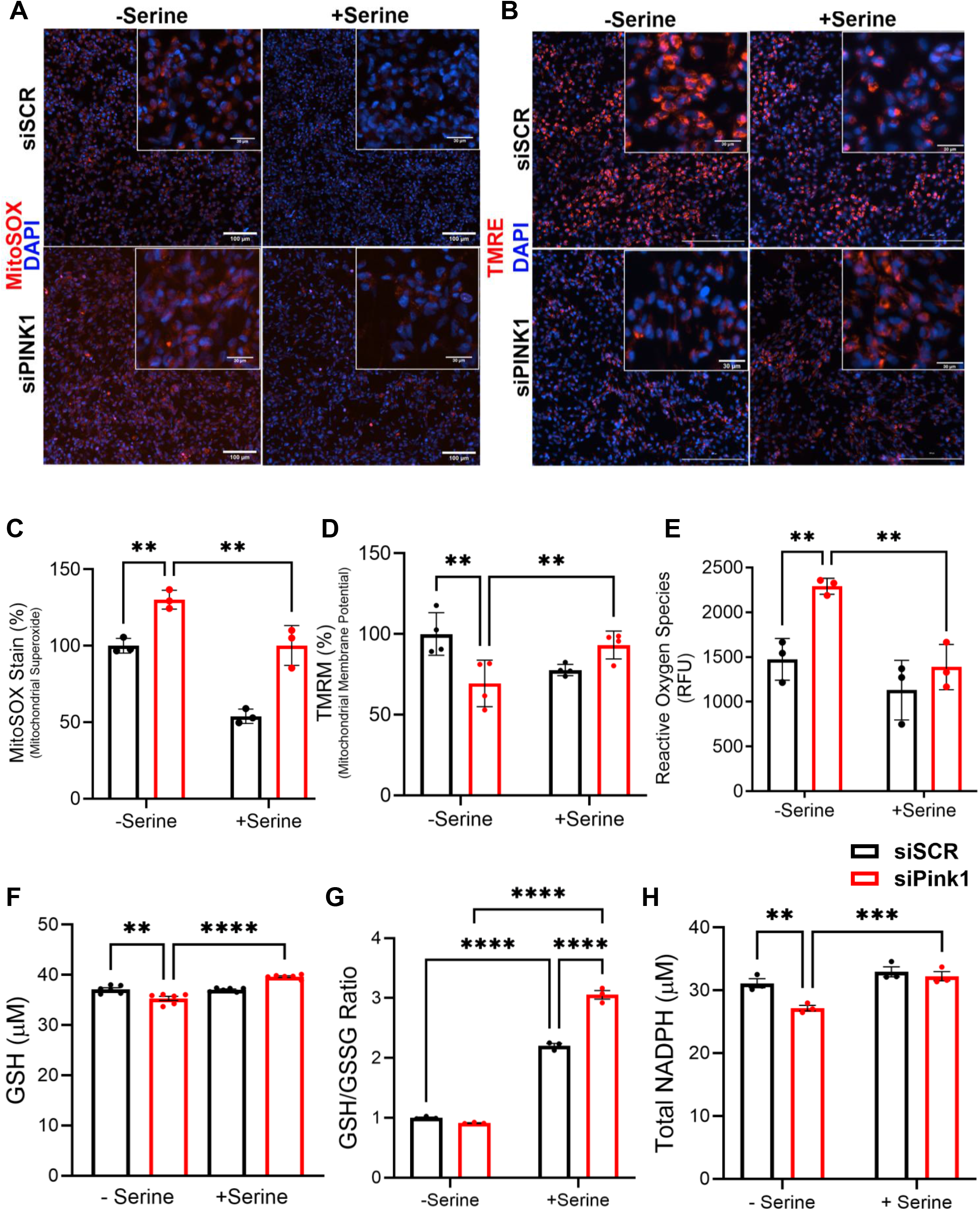
Serine supplementation rescues mitochondrial homeostasis and ROS management systems. ARPE-19 cells were treated with 20nmols siPink1 for 5 days and serine was supplemented at 200uM refreshed at 3 days. (**A, C**) siPink1 ARPE-19 cells were treated with 1uM MitoSOX Red fluorescent stain (Sigma M36007) for 30 minutes and nuclei were counterstained with 1ug/mL Hoechst 33342. (**B, D**) Mitochondrial membrane potential was measured with TMRM (Sigma T5428). siPink1 treated cells were incubated with 1uM TMRM for 30 minutes followed by nuclear stain with 1ug/mL Hoechst 33342. (**E**) Total cellular ROS was measured using DHE based fluorescence kit (Cayman 601290). (**F, G**) Total Glutathione and GSH/GSSG ratios were assessed with a redox based colorimetric kit (Abcam ab138881). (**H**) Total NADPH levels were measured using colorimetric NADP+/NADPH kit (Abcam ab186033). Data are presented as mean +/- SEM from n = 3 replicates per group. Statistical significance was determined using two-way ANOVA with siPINK1 (knockdown vs. control) and +/- serine supplementation, followed by Tukey’s post hoc multiple comparisons test. (**p<0.005; ***p<0.0005; ****p<0.0001)

### PINK1 loss dysregulates AMPK-mTOR-4EBP1 signaling

We set to investigate how Pink1 loss and serine supplementation modulates nutrient stress and its downstream stress response signaling. Phosphorylation states of AMPK, AKT, mTOR, 4EBP1, and ATF4 were assessed by Western blot in siPINK1-transfected ARPE-19 cells ± 200 µM serine supplementation for 5 days. AMPK phosphorylation at T172 was elevated by 1.5-fold in siPINK1 cells and was reduced back to basal levels by serine supplementation **(Fig. 4A, F)**. p-AKT (S473) was modestly elevated in siPINK1 cells and reduced by serine supplementation. **(Fig. 4B)**. Next, we tested the involvement of Pink1 loss on mTOR by measuring p-mTOR levels which were increased siPINK1 cells by 1.5-fold and was rescued back to basal levels upon serine supplementation **(Fig. 4C,I)**. Next, we assessed the downstream translation activator, 4EBP1. p-4EBP1 was increased in siPINK1 cells by 1.4-fold and further elevated to 1.7-fold with serine supplementation **(Fig. 4D, H)**. ATF4 mRNA was elevated by 1.4-fold in siPINK1 cells **(Fig. S2F)**; however, ATF4 protein was reduced by 30% and was not restored by serine supplementation **(Fig. 4E, J)**. In PINK1^−/−^ mouse RPE, ATF4 mRNA was significantly reduced relative to wild-type and was rescued by 6 months of dietary serine supplementation. These data suggest that the AMPK/mTOR energy sensing signaling pathways are altered with Pink1 loss and serine supplementation can restore AMPK and mTOR signaling pathways.

**Figure 4.**
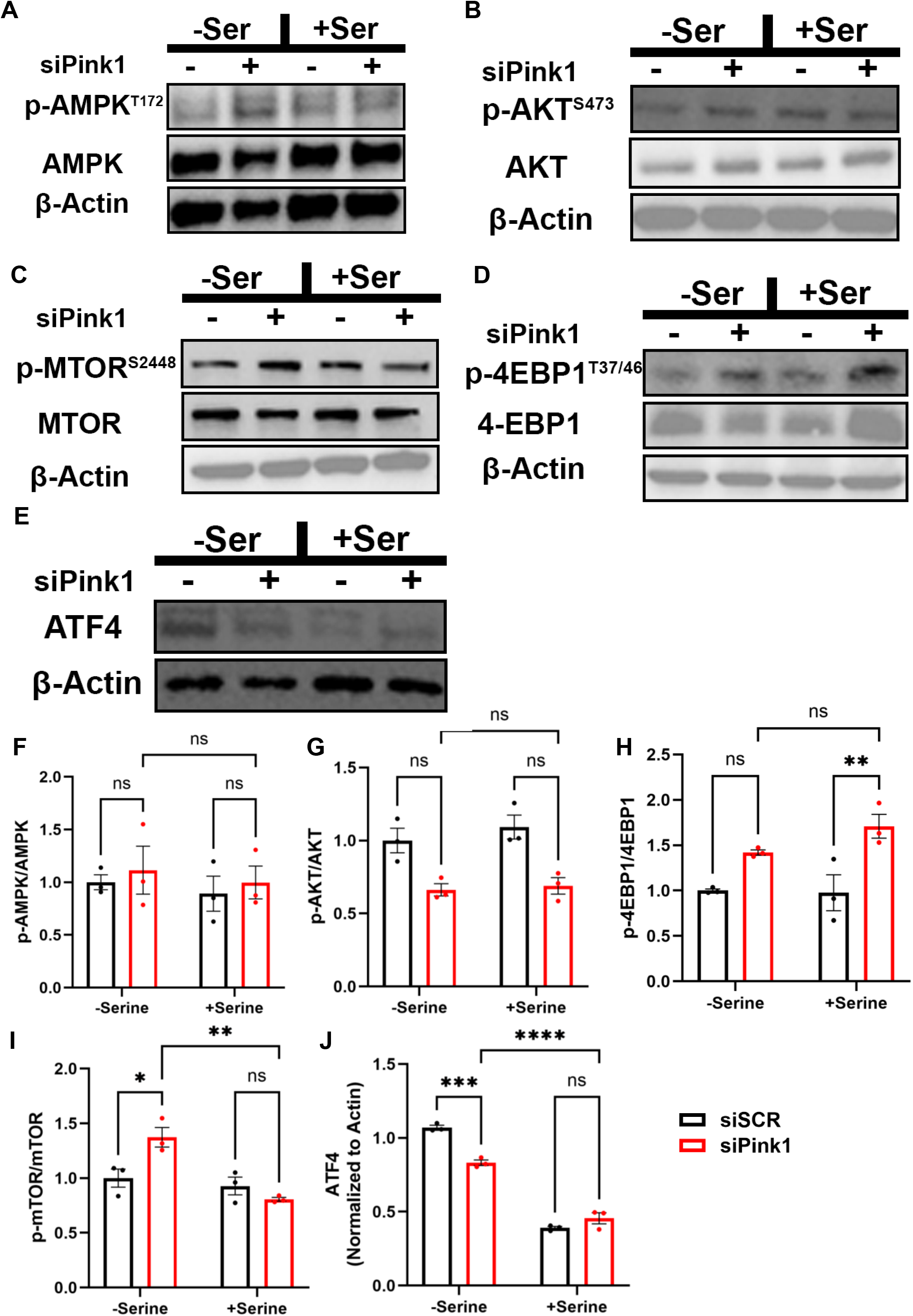
Pink1 loss alters AMPK, mTOR and ATF4 expression. ARPE-19 cells were treated with 20nmols siPink1 for 5 days. Cells were supplemented with 200uM serine refreshed at 3 days. Cells were pelleted and lysed in RIPA buffer and 10ug was loaded for immunoblotting for the following proteins. (**A, F**) p-AMPK (T172) (CST2535S) and total AMPK (CST5831S) (**B, G**) p-AKT (S473) (CST12694) and total AKT (CST4691) (**C, H**) p-4EBP1 (Thr 37/46) (CST9451S) and total 4EBP1 (CST9452). (**D, I**) p-MTOR (Ser2448) (CST2971) and total mTOR (CST2971) (**E, J**) ATF4 (CST11815) (**F**). Representative blots from n = 3 replicates per group are shown. Statistical significance was determined using two-way ANOVA with siPINK1 (knockdown vs. control) and +/- serine supplementation, followed by Tukey’s post hoc multiple comparisons test. (*p<0.05; **p<0.005;***p<0.0005)

### PINK1 deficiency alters inflammatory profile of the RPE and partially mitigated by serine supplementation

Next, we examined the potential involvement of Pink1 in regulating the inflammatory response elements within the RPE. IL-6 was upregulated in siPINK1 cells by ∼1.8-fold, however, was not reduced by serine supplementation (**Fig. 5A**). IFI6 expression was unchanged by PINK1 loss but was significantly reduced by serine supplementation (**Fig. 5B**). STAT3 expression was unchanged by PINK1 knockdown but was markedly reduced by serine supplementation in both siScramble and siPINK1 cells (**Fig. 5C**). FOXO1 was reduced in serine treated cells regardless of Pink1 loss. Surprisingly, FOXO3 was significantly elevated by ∼4-fold in siPINK1 cells and like FOXO1, serine supplementation reduced its expression in both siScramble and siPINK1 conditions (**Fig. 5D, E**). Next, We assessed the status of NF-κB activation via p65 phosphorylation (Ser536) which was unchanged across all conditions (**Fig. 5F, top panel**). NLRP3 protein levels were significantly elevated in siPINK1 cells and were not reduced by serine supplementation (**Fig. 5F, bottom panel**). These data suggest that serine is involved in the regulation of several interlinked inflammatory mediators.

**Figure 5.**
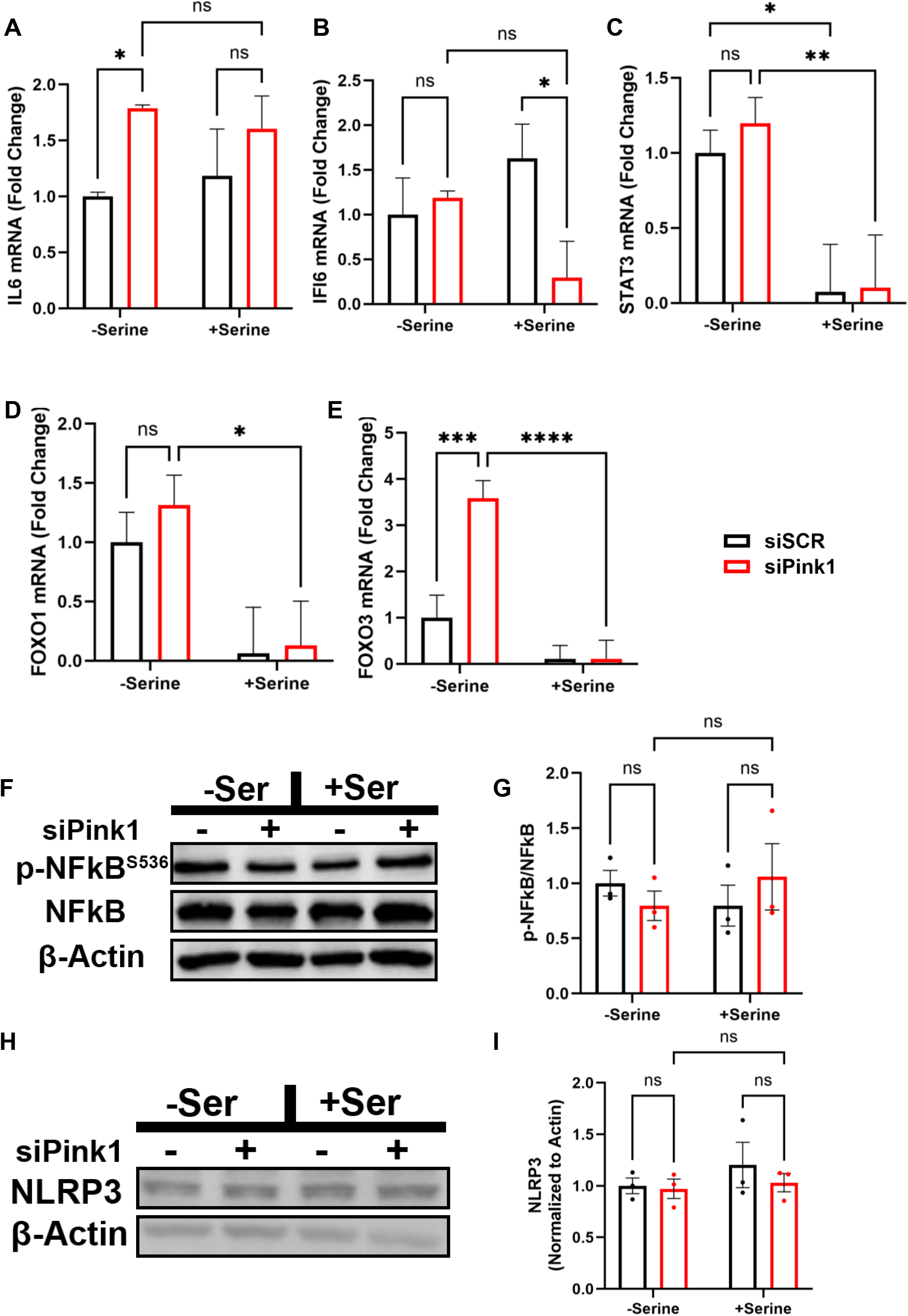
PINK1 deficiency alters inflammatory profile of the RPE and partially mitigated by serine supplementation. ARPE-19 cells were treated siPink1 and were supplemented with 200uM serine refreshed at 3 days. RNA was extracted from these cells using ZYMO RNA kit (R1058). qPCR was performed on a BioRad Opus thermocycler for (**A**) FOXO1 (**B**) FOXO3IFI6 (**C**) IL6 (**D**) IFI6 and (**E**) STAT3. 10ug of ARPE-19 protein lysate was used for immunoblotting for: (**F, G**) p-NFkB (Ser536) (MAB7226). (**H, I**) NLRP3 (NBP2-12446) activation was increased in siPink1 treated ARPE-19 cells and serine supplementation reduced NLRP3 activation. Representative blots from n = 3 replicates per group are shown. Statistical significance was determined using two-way ANOVA with siPINK1 (knockdown vs. control) and +/- serine supplementation, followed by Tukey’s post hoc multiple comparisons test. (*p<0.05; **p<0.005; ***p<0.0005; ****p<0.0001)

### Serine supplementation rescues *in vivo* retinal structure and function

To assess retinal and RPE structures *in vivo*, we examined color fundus structures, OCT images and BlamD deposits. Following this, we assessed retinal function via electroretinogram (ERG) and tested optokinetic responses (OKR). WT and PINK1^−/−^ mice were aged 18 months and administered 0.5% w/v L-serine in drinking water ad libitum for 6 months (**Fig. 6A**). L-serine levels in RPE lysates were significantly reduced in PINK1^−/−^ mice by ∼40% relative to wild-type controls (**Fig. 6B**). Retinal cross-sections stained with H&E revealed disorganized retinal architecture in PINK1^−/−^ mice, including outer nuclear layer thinning and irregular inner and outer segment spacing (**Fig. 6C**). PHGDH immunostaining was significantly reduced in the RPE monolayer and outer nuclear layer of PINK1^−/−^ mice and was restored by serine supplementation, which also normalized RPE serine concentrations to near wild-type levels (**Fig. 6D**). At the mRNA level, PINK1 and PHGDH transcripts were both reduced in PINK1^−/−^ RPE lysates and were increased by serine supplementation (**Fig. S4A**). Expression of other serine synthesis pathway genes was not significantly altered by PINK1 loss or serine supplementation (**Fig. S4B-G**). ZO-1 flatmount staining revealed disrupted RPE morphology in PINK1^−/−^ mice, with irregular cell shapes, fragmented junctions, and loss of hexagonal mosaic organization (**Fig. 6E**); these features were largely restored in serine-supplemented PINK1^−/−^ mice. Transmission electron microscopy revealed swollen mitochondria with disrupted cristae in PINK1^−/−^ RPE, which were restored to normal morphology with tightly packed cristae following serine supplementation (**Fig. 6F**).

**Figure 6.**
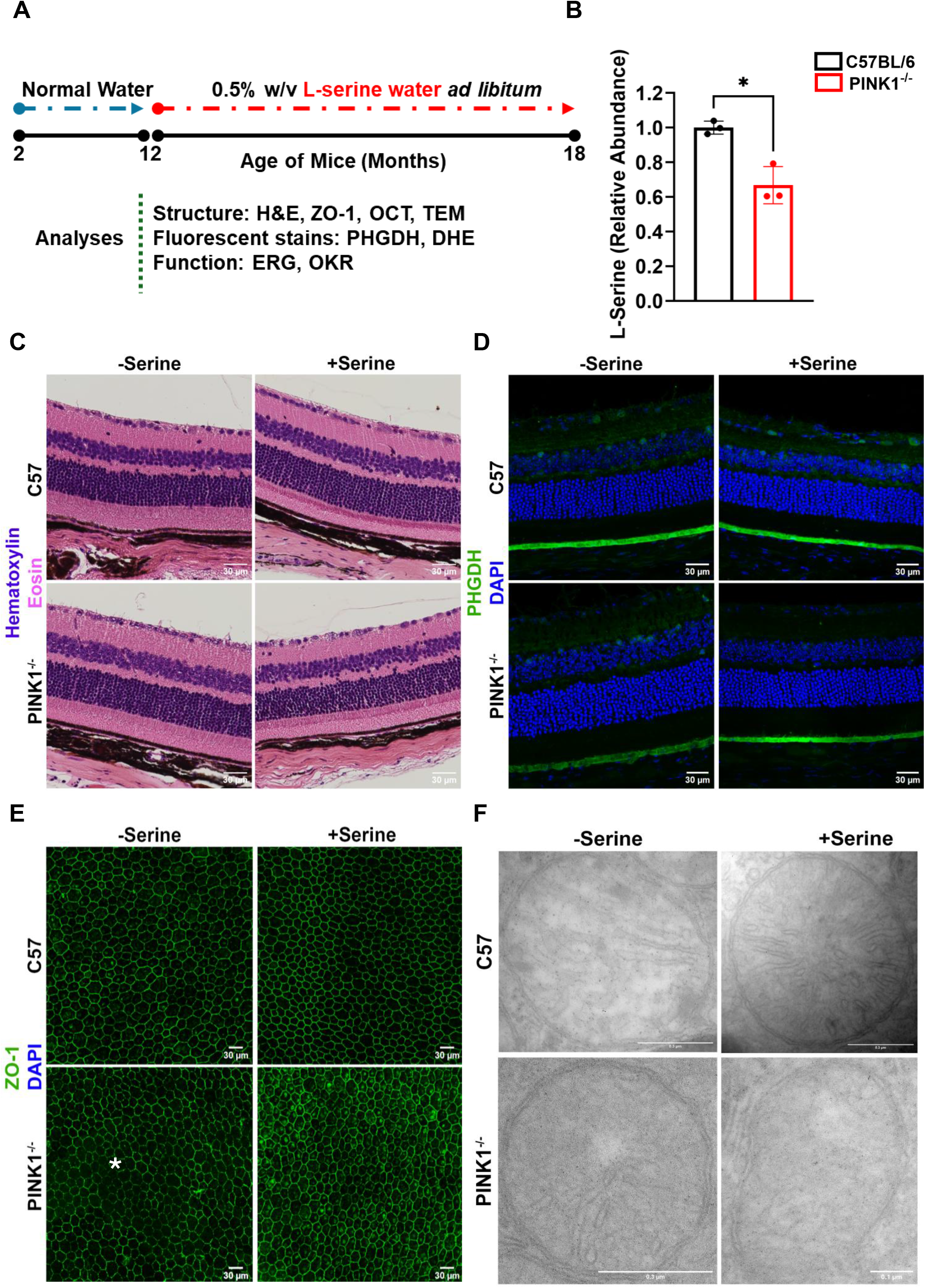
Serine supplementation alters RPE and mitochondrial morphology. (**A**) C57BL/6 and PINK1^-/-^ mice were aged to 18 months followed by 24 weeks of *ad libitum* 0.5% w/v serine in drinking water. (**B**) Serine abundance in the isolated RPE layer was analyzed using serine kit (MAK352). Formalin fixed paraffin embedded and sectioned eyes were used and images were captured from the peripheral retina. (**C**) Hematoxylin and Eosin stains for retinal structures (**D**) Immunofluorescent stain for PHGDH (Sigma HPA021241). (**E**) Retinal flatmounts were processed for ZO-1 immunofluorescence (SCBT-sc33725 AF488) and images were captured from the peripheral RPE. White star marks region of dysmorphic RPE in Pink1-/- mice. (**F**) Transmission electron microscopy (TEM) was performed on ultra-thin sections and were imaged at 60kV and 48Kx magnification. Representative images from n = 6 replicates per group are shown. Statistical significance was determined using student t-test for C57BL/6 vs Pink1^-/-^. (*p<0.005)

Furthermore, *in vivo* DHE staining revealed elevated superoxide signals in the RPE and photoreceptor outer segment layers of PINK1^−/−^ mice, which was significantly reduced by serine supplementation (**Fig. 7A**). GFAP immunostaining was reduced in PINK1^−/−^ retinas relative to WT and was partially restored by serine supplementation (**Fig. 7B**). IBA1 immunostaining revealed increased microglial activation in the inner and outer plexiform layers of PINK1^−/−^ retinas, which was reduced in serine-supplemented PINK1^−/−^ mice (**Fig. 7C**). Circulating IL-6 levels were elevated in serum from PINK1^−/−^ mice, however, dietary serine supplementation did not rescue IL-6 elevation (**Fig. 7D, E**). These data combined with *in vitro* inflammatory markers suggest that Pink1 is involved in the regulation of pro-inflammatory mediators in the RPE, and serine supplementation could lead to a reduction in overall inflammatory load.

**Figure 7.**
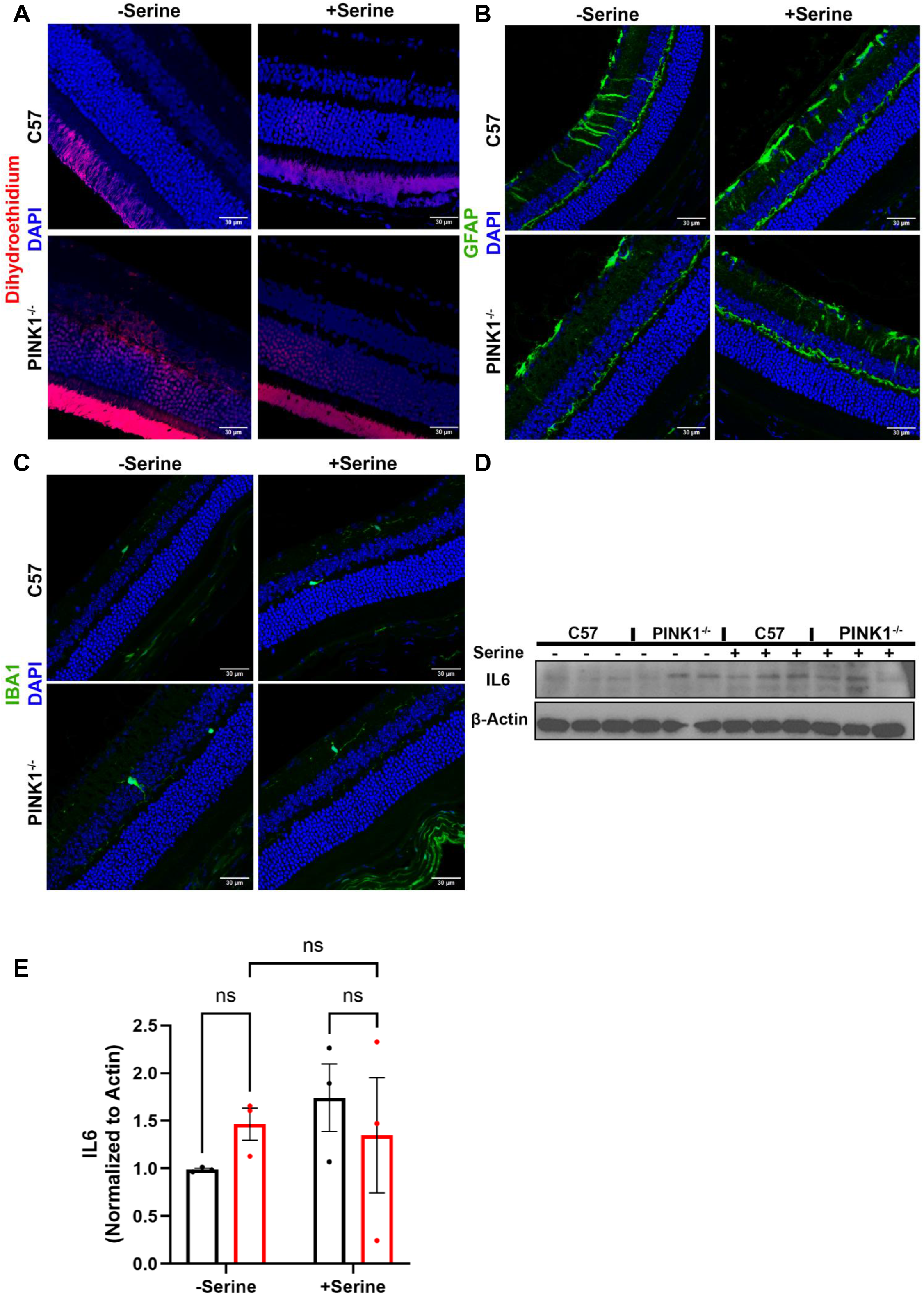
Serine supplementation reduces superoxides and improves Muller cell activation. C57BL/6 and PINK1-/- mice supplemented with either normal water or serine water for 24 weeks. Paraffin embedded tissues were used for the following fluorescent imaging. Sections were imaged at the peripheral retina. (**A**) Dihydroethidium (DHE) stain for ROS mediated damage. (**B**) Muller cell activation was recorded with GFAP activation within the retina and the retinal ganglion cell layer. (**C**) Microglial activation via IBA1. (**D, E**) Western blot from serum samples of mice from each group were probed for circulating pro-inflammatory IL6 levels (abcam ab229381). Immunostains and western blot were performed in eyes from n=3 mice and representative images are reported.

Fundus imaging revealed no gross structural differences across groups (**Fig. 8A**). Fundus autofluorescence imaging revealed hyper-reflective foci in PINK1^−/−^ mice that were markedly reduced by serine supplementation (**Fig. S4H**). OCT quantification showed increased photoreceptor and total retinal thickness in PINK1^−/−^ mice, which were reduced by dietary serine (**Fig. 8B, C**). TEM imaging revealed BLamD-like deposits between the RPE basal membrane and Bruch’s membrane in PINK1^−/−^ mice that were largely absent in WT and serine-treated animals (**Fig. 8D**). Full-field ERG recordings in 12-hour dark-adapted mice showed significantly reduced a-, b-, and c-wave amplitudes in PINK1^−/−^ mice; serine supplementation significantly increased a- and b-wave amplitudes and c-wave amplitude and extended c-wave implicit time (**Fig. 8E–G**). Optokinetic responses were not significantly different across groups (**Fig. S4I**). These data suggest that functional deficits brought about by the loss of Pink1 can be rescued by dietary serine supplementation. Together, these data establish that PINK1 deficiency disrupts serine biosynthesis, mitochondrial redox balance, and induces RPE dysmorphology, and that dietary serine supplementation serine rescues oxidative stress, inflammation and RPE and retinal dysfunction.

**Figure 8.**
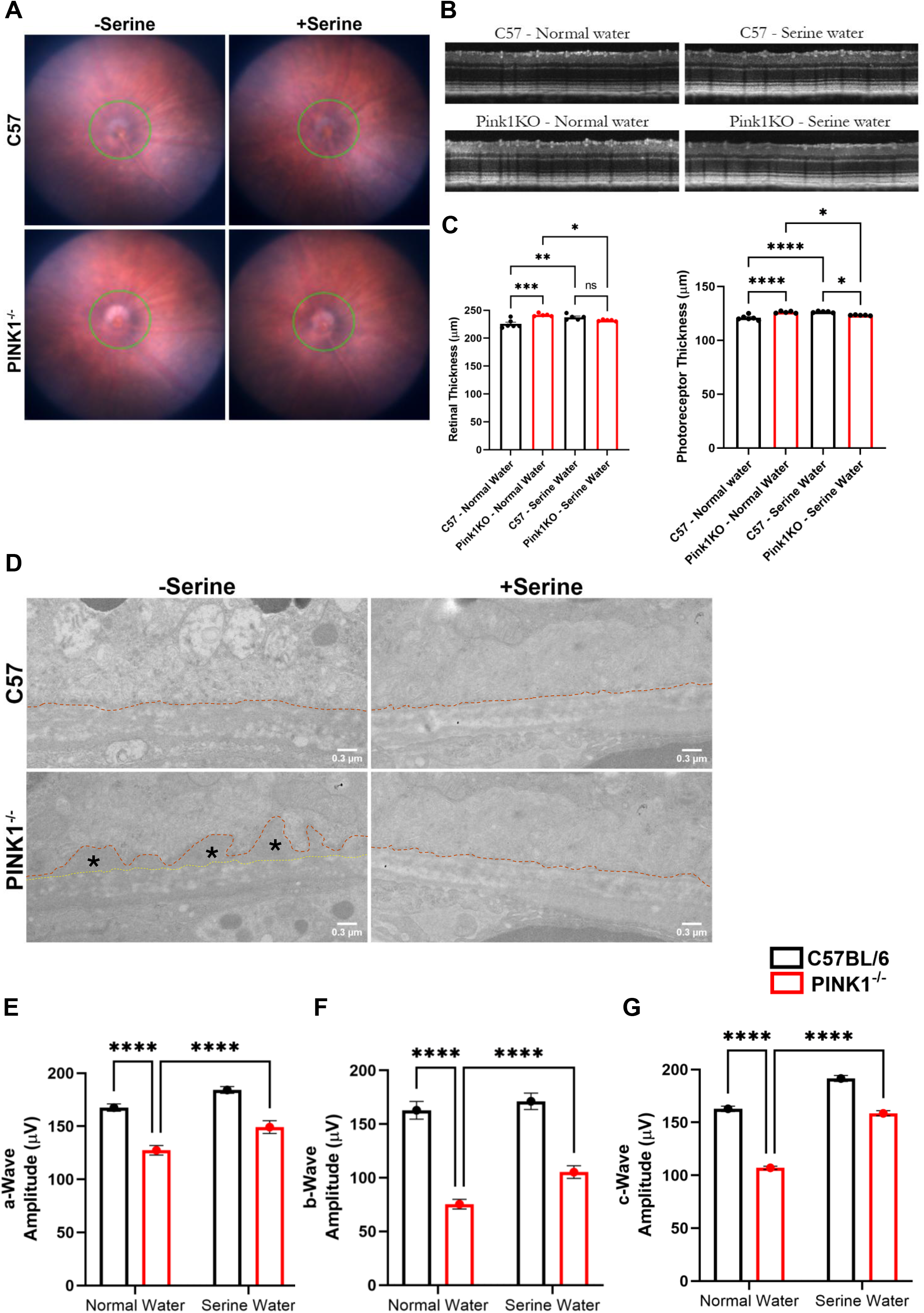
Serine supplementation rescues retinal structures and function. C57BL/6 and PINK1-/- mice supplemented with either normal water or serine water for 24 weeks. (**A**) Color fundus images were acquired using the MICRON IV system. (**B**) OCT module of the Micron IV system was used to capture segments adjacent to the optic nerve. (**C**) Retinal and photoreceptor thicknesses were measured using Photoshop pixel counter (**D**) TEM imaging of the retina was performed on ultra-thin sections and were imaged at 60kV and 4.8Kx magnification. The RPE layer was identified, and basal laminar deposits were marked with (*). Electroretinography (ERG) responses from dark adapted mice were recorded. (**E**) Photoreceptor responses to light stimulus were recorded as the a-wave amplitude. (**F**) ON bipolar cells and Muller cell responses were recorded as b-wave amplitude. (**G**) RPE layer’s responses were recorded as the c-wave amplitude. Each measurement was recorded with n=6 animals with averages of both eyes. Statistical significance was determined using two-way ANOVA with PINK1 (knockout vs. C57) and +/- serine supplementation, followed by Tukey’s post hoc multiple comparisons test. (*p<0.05; **p<0.005; ***p<0.0005; ****p<0.0001)

## Discussion

Age-related macular degeneration is characterized by the aberrant accumulation of ROS leading to mitochondrial impairment, switch to glycolysis over mitochondrial oxidative phosphorylation, chronic inflammation and progresses into RPE dysfunction^3-5,8,9^. Here, we show that PINK1 is involved in the regulation of serine metabolism which can directly influence the RPE’s capacity to combat reactive oxygen species. In AMD patient tissues, we show localized loss of PHGDH in the RPE which suggests that a localized serine synthesis vulnerability exists and could lead to AMD pathophysiology driving RPE dysfunction. In both ARPE-19 cells and PINK1^−/−^ mice, serine supplementation restored metabolic balance, attenuated oxidative stress, and preserved RPE and retinal integrity, establishing a mechanistic link between mitochondrial quality control and one-carbon metabolism with therapeutic relevance for AMD.

PINK1 loss reduced PHGDH expression and intracellular serine levels, consistent with impaired de novo serine synthesis. GSH levels, the GSH/GSSG ratio, and NADPH pools were all reduced in siPINK1 cells, accompanied by elevated cellular and mitochondrial ROS. Direct PHGDH knockdown recapitulated these redox defects, confirming that serine metabolism mediates the observed phenotype. Serine supplementation restored GSH and NADPH levels and suppressed ROS accumulation. Notably, exogenous serine did not restore PHGDH transcript or protein levels *in vitro*, indicating that the biosynthetic deficit persists despite replenishment of the serine pool. In contrast, dietary serine supplementation in PINK1^−/−^ mice increased both PINK1 and PHGDH mRNA, suggesting a positive feedback relationship between serine availability and pathway gene expression, possibly via ATF4 modulation.

Mitochondrial membrane potential was reduced, and mitochondrial superoxide was elevated in siPINK1 cells; both were improved by serine supplementation, consistent with enhanced antioxidant capacity indirectly stabilizing mitochondrial integrity. TEM analysis of PINK1^−/−^ RPE revealed distorted cristae and sub-RPE deposits, both of which were ameliorated by dietary serine. Importantly, serine supplementation did not rescue basal or maximal oxygen consumption rate, indicating that mitochondrial respiratory capacity remains impaired despite improvements in redox balance and membrane potential. These findings suggest that serine acts directly to augment the antioxidant deficit and improve mitochondrial ROS quenching but does not intervene with oxidative phosphorylation.

PINK1 knockdown elevated AMPK phosphorylation, modestly increased p-AKT, and increased p-mTOR, reflecting a state of concurrent energetic stress and dysregulated anabolic signaling. Serine supplementation reduced p-AMPK, p-AKT, and p-mTOR. However, p-4EBP1 was paradoxically elevated by serine supplementation, suggesting that serine may selectively redirect mTORC1 translational outputs rather than uniformly suppressing mTORC1 activity. ATF4 levels were reduced in siPINK1 cells despite elevated ATF4 mRNA, pointing to post-transcriptional suppression of integrated stress response output and serine supplementation did not restore ATF4 protein *in vitro*. In contrast, PINK1^−/−^ mouse RPE showed a significant decrease in ATF4 mRNA and was rescued by dietary serine. This highlights the possibility of long-term serine supplementation can modulate *in vivo* ATF4 abundance thereby aiding in the integrated stress response pathway.

FOXO1 and FOXO3 were elevated in siPINK1 cells and suppressed by serine supplementation in both knockdown and control conditions. Among downstream inflammatory markers, IL-6 was elevated by PINK1 loss but was not reduced by serine supplementation in vitro, whereas IFI6 and STAT3 were suppressed by serine independently of PINK1 status. NF-κB phosphorylation was unchanged across all conditions. NLRP3 was elevated in siPINK1 cells and was not reduced by serine supplementation, indicating that inflammasome activation is independent of serine availability. PINK1^−/−^ mice displayed elevated serum IL-6, increased Iba1-positive microglial activation, and reduced GFAP, all of which were improved by dietary serine, suggesting that serine is tightly integrated with the systemic and retinal inflammatory responses as observed in other inflammatory mouse models^13^.

PINK1 loss produced structural deterioration in the mouse RPE and retina, including outer nuclear layer thinning and disorganized retinal lamination on H&E, disrupted ZO-1 continuity and dysmorphic RPE morphology on flatmounts, BLamD-like sub-RPE deposits on TEM, and increased photoreceptor and total retinal thickness on OCT. Fundus autofluorescence revealed hyper-reflective foci in PINK1^−/−^ mice. Dietary serine supplementation ameliorated these structural abnormalities. Full-field ERG demonstrated significantly reduced a-, b-, and c-wave amplitudes in PINK1^−/−^ mice, with serine supplementation rescuing all three parameters. However, serine supplementation had no effect on contrast sensitivity.

Collectively, these data support a model in which PINK1 deficiency imposes dual metabolic stress: impaired mitophagy drives ROS accumulation, while reduced PHGDH-dependent serine synthesis limits redox buffering capacity. Exogenous serine bypasses the biosynthetic deficit, replenishing GSH and NADPH thereby suppressing oxidative stress, and rescuing RPE and retinal structure and function. Recent studies in patients with MacTel Type 2 show that loss of PHGDH and serine levels in the retina is implicated in disease pathophysiology^16,19,20^ and serine supplementation is currently being investigated in clinical trials (ClinicalTrials.gov NCT04907084). These studies mainly show that serine is safe at high doses, and dietary serine readily enters the retina. Therefore, a similar serine supplementation regimen could be a viable strategy to supplement the current standard of AREDS2 therapy for AMD.

## Materials and Methods

### Cell Culture

AMD patient derived induced pluripotent stem cell-derived retinal pigment epithelial (iPSC-RPE) cells and ARPE-19 cells were used in all experiments. ARPE-19 cells were maintained in DMEM/F12 media (Thermo Fisher Scientific) supplemented with 10% fetal bovine serum (FBS), at 37°C in a 5% CO_2_ humidified incubator.

### iPSC-RPE differentiation

Human AMD donor induced pluripotent stem cells (iPSCs) from non-AMD, early AMD and advanced AMD patients along with IMR90-4 controls were differentiated into RPE using previously established protocols. Briefly, iPSC were seeded onto vitronectin coated 6 well plates in E8 medium. They were then switched to retinal initiation media (RIM) for two days. They were then switched to retinal differentiation media (RDM) until day 14 followed by retinal differentiation media supplemented with 10mM nicotinamide (RMNA) until day 25. The cells were then switched to RPE maturation media (RPE-MM) until day 75. Over the first ∼2–4 weeks, cells underwent neuroectodermal and retinal lineage specification, forming pigmented islands characteristic of RPE commitment. Pigmented epithelial clusters were manually enriched or isolated using gentle dissociation and replated to promote monolayer formation and maturation. Once expanded, cultures were maintained in RPE maturation medium, and cells were cultured for an additional 4–8+ weeks until they exhibited cobblestone morphology, uniform pigmentation, and a stable monolayer. Daily to every-other-day medium changes were performed throughout differentiation and maturation.

Mature iPSC-RPE cultures were validated by assessing expression of stem markers (TUBB3, AFP and RBFOX3) and canonical RPE markers (MITF, BEST1, RPE65, ZO-1) via immunocytochemistry, qPCR and western blots.

### Serine/Glycine Free Medium

Custom amino acid media was made from MEM (ThermoFisher 11095080) with the following formulation **(Table 1)**, which does not contain serine for serine deprivation experiments. To make serine/glycine-replete medium, L-Serine (0.4 mM) and L-Glycine (0.4 mM) were added to serine/glycine free medium.

**Table 1.**
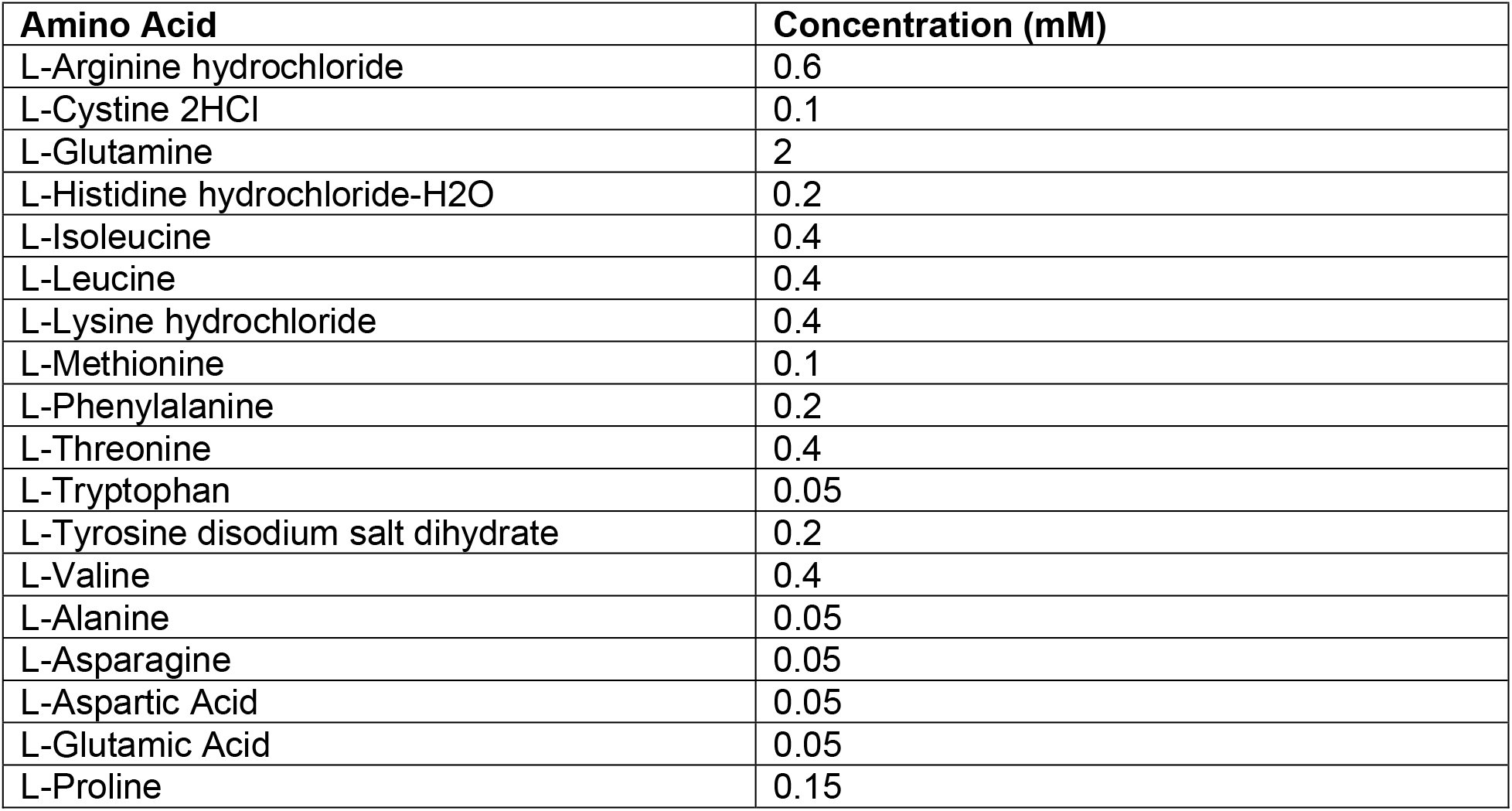
Custom amino acid formulation.

**Table 2.1.**
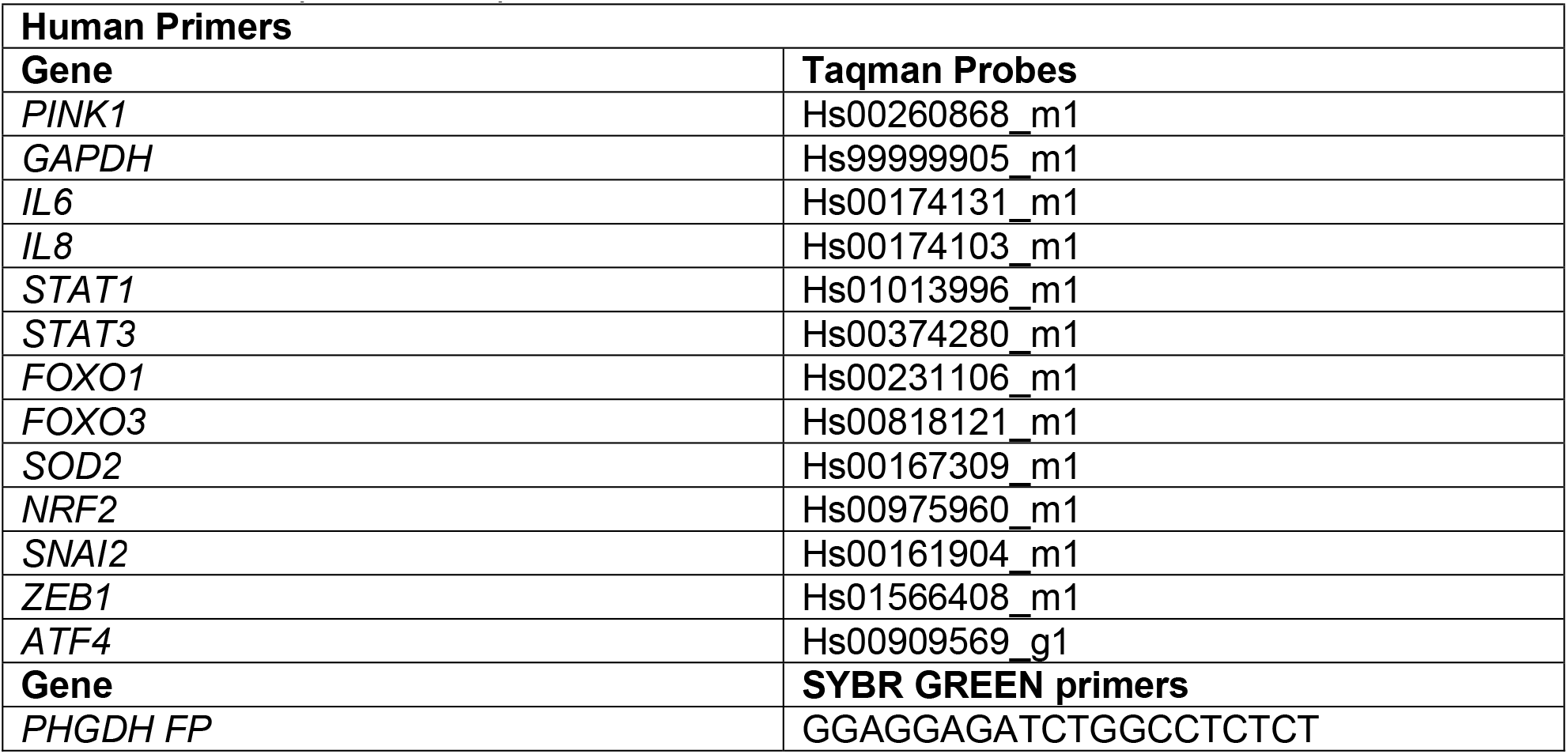
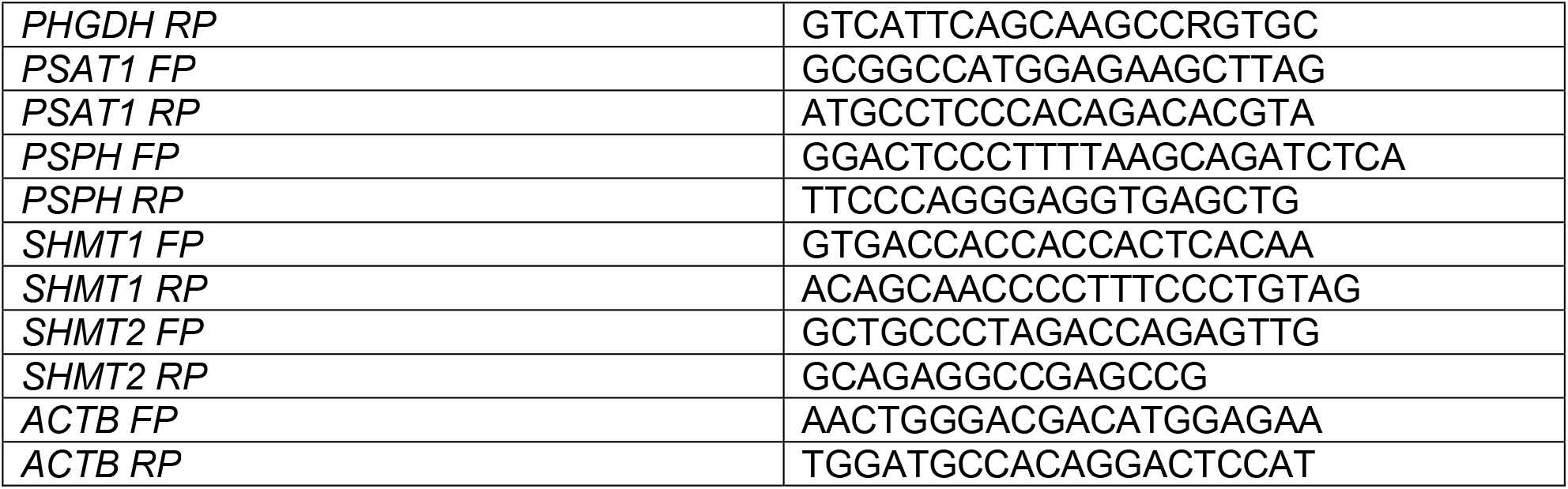
Human primers for qPCR.

**Table 2.2.**
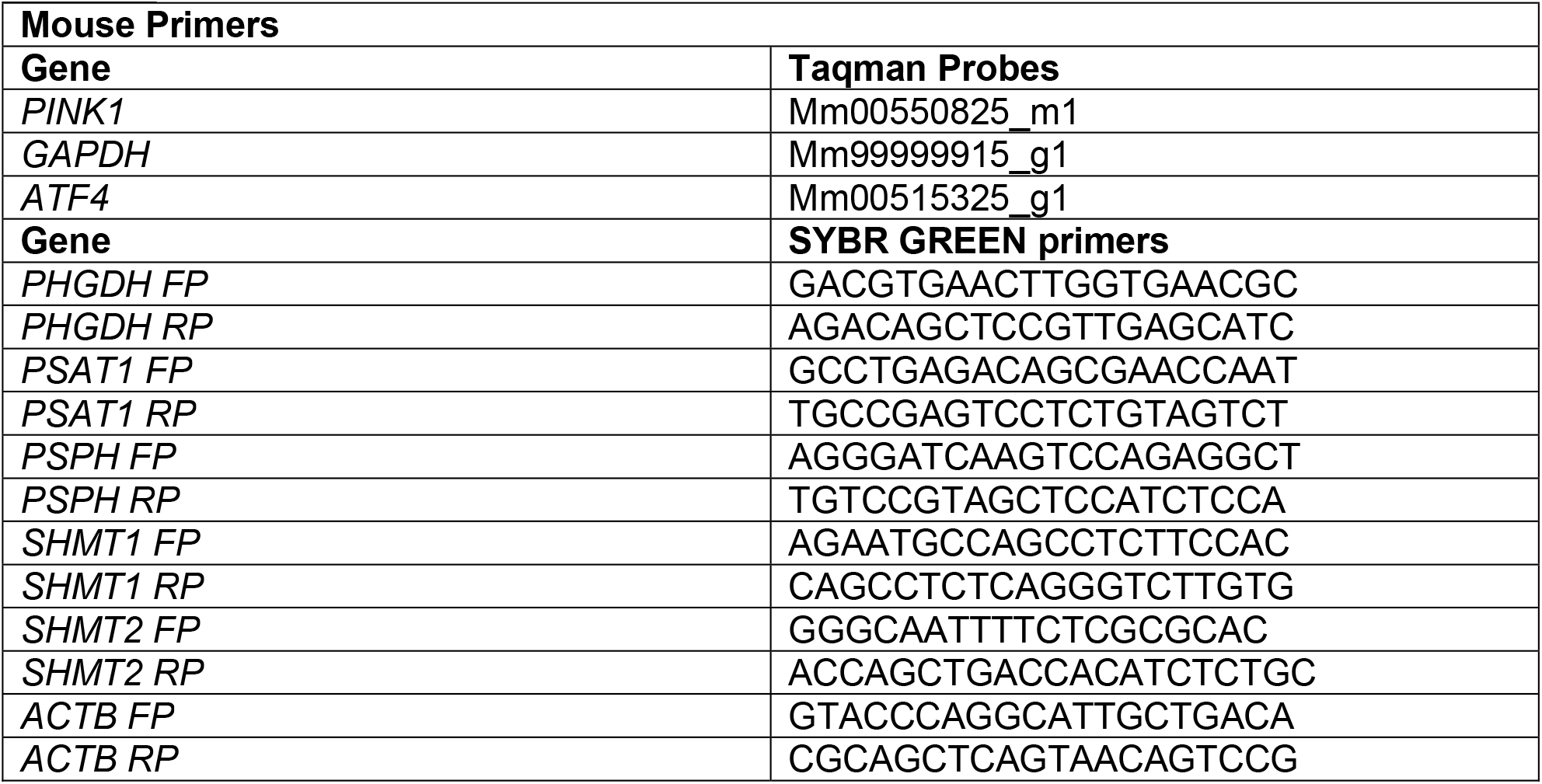
Mouse primers for qPCR.

### siRNA treatments

For knockdown experiments, ARPE-19 cells were transfected with 20pmol human PINK1 siRNA (Thermo AM51331) or 20pmol mouse PINK1 siRNA (Thermo AM16708) using 3.5ug/well Lipofectamine 2000 (Thermo 11668019) for 5 days.

### L-Serine supplementation

ARPE-19 cells were treated with 200 µM L-Serine for 5 days with a refresh on day 3 to assess rescue effects. Serine free media was used as control for serine replete media.

### GSH and GSSG Measurement

ARPE-19 cells treated with siPink1 +/- serine were processed according to manufacturer instructions (Abcam ab138881). Briefly, GSH and GSSG standards (0.1563-10uM) were prepared. 2x10^6^ cells were lysed in lysis buffer and deproteinized in tricholoroacetic acid and neutralized with NaOH (pH 7.4). They were then centrifuged at 10,000g for 15 minutes at 4C. GSH and GSSG detection probes were then added and fluorescence λex = 490 nm and λem = 520 nm. was measured at 15-minute intervals for 1 hour.

### NADPH and NADH measurements

ARPE-19 cells were lysed and processed according to manufacturer instructions (Abcam 186031). 2x106 cells were lysed in lysis buffer at room temperature for 15 minutes. The samples were then centrifuged at 2500 rpm for 10 minutes. The supernatant was collected and 50uL was added to a black walled 96 well plate and incubated with the provided NADPH reaction mixture. Absorbance was measured at 460nm at 15-minute intervals for a period of 1 hour.

### RNA isolation

Total RNA was isolated from cultured ARPE-19 cells and mouse RPE explants and iPSC-RPE cells using Zymo Research Miniprep plus (R1058). RNA from RPE layer was isolated as previously described. Briefly, eyes were enucleated and the anterior chamber was removed. Retina was gently separated and the underlying RPE layer was cut out into petals and placed in a well of a 96 well plate. 100uL of RNA lysis buffer was gently layered on. The plate was shaken for 15 minutes, and the supernatant was collected and RNA was isolated. 1ug of RNA was converted into cDNA using Fisher Scientific 43-688-14.

### qPCR

qPCR for the genes listed below was performed using Taqman and SYBR green (Neb 43-688-14).

### Immunoblotting

Cell lysates were prepared in RIPA lysis buffer with protease and phosphatase inhibitor cocktail. Mice were euthanized and the RPE layer was collected as previously described<ref>. Eyes were enucleated and the retinas were recovered. The underlying RPE layer was cut into petals, transferred to a 96 well plate and was layered with RIPA buffer on the RPE for 15 minutes. Protein concentration was measured using DC assay (Biorad 5000111). 10ug of supernatant lysate was subjected to western blotting. SDS PAGE was performed on 10% or 12% gels and transferred onto nitrocellulose membranes. Membranes were blocked for one hour in 5% non-fat dry milk at room temperature. Primary antibodies (Table 2) were diluted at 1:1000 and were incubated overnight at 4C. Membranes were washed with 1x TBS and 0.1% Tween-20 and detected with Enhanced Chemiluminescence (ECL) substrate (Thermo Fisher 32106).

**Table 2.** Antibodies used for Immunofluorescence and Western blots.

| **Antibody** | **Cat No:** |
| --- | --- |
| PHGDH | CST13428 |
| Pink1 | SCBT sc-33796 |
| β-Actin | SCBT sc-47778 |
| E-Cadherin | SCBT sc-8426 |
| Vimentin | SCBT sc-373717 |
| p-AMPK | CST2535S |
| AMPK | CST5831S |
| p-MTOR | CST2971 |
| MTOR | CST2972 |
| p-4EBP1 | CST9451S |
| 4EBP1 | CST9452 |
| p-AKT | CST12694 |
| AKT | CST4691 |
| p-NFκB | MAB7226 |
| NFκB | AF5078 |
| NLRP3 | NBP2-12446 |
| IL6 | Ab233706 |
| ATF4 | CST11815 |
| ZO-1 | SCBT-sc33725 AF488 |
| GFAP | DAKO Z0334 |
| IBA1 | ThermoFisherMA5-37992 |
| AlexaFluor 568 | ThermoFisher A10042 |
| AlexaFluor 647 | ThermoFisher A32795 |

Antibodies used are listed in Table1. Goat anti Rb (ab205718) and Rabbit anti Ms (ab6728) secondary antibodies were used at 1:2000.

### Brightfield Microscopy

Brightfield images were acquired using EVOS Imager at 4x, 10x and 20x as indicated.

### Immunofluorescence

For immunofluorescence imaging, eyes were enucleated and fixed in a 97% methanol:3% glacial acetic acid solution for 1 week. Eyes were then processed through gradient alcohols and embedded in paraffin blocks. These blocks were sectioned at 5um and were dried overnight. Slides were rehydrated through xylenes and gradient alcohols and PBS. Antigen retrieval was performed in 10mM citrate buffer (pH 6.0) for 20 minutes. Slides were washed with PBS and blocked in 1% BSA solution for 1 hour. Primary antibodies were diluted in 1% BSA in PBS and incubated overnight at 4°C. After washing with PBS with 0.01% tween-20, sections were incubated with AlexaFluor 568 (A10042) or AlexaFluor 647 (A32795) secondary antibodies (1:500, Thermo Fisher Scientific) for 1 hour at room temperature. Nuclei were counterstained with DAPI (1µg/mL) for 5 minutes. Coverslips were mounted on glass slides with ProLong Gold Antifade reagent. Images were captured using NIKON AX confocal microscope.

### Retinal Flatmounts

Retinas were harvested from euthanized mice, fixed in 4% paraformaldehyde (PFA) for 1 hour at 4°C, and dissected to remove the cornea, lens, and sclera, leaving the retina. Flatmounts were prepared by making radial cuts to flatten the retina, followed by permeabilization in 0.1% Triton X-100 for 30 minutes, incubated in 5% BSA for 1 hour at room temperature, then with ZO-1 primary antibody (1:200, Santa Cruz sc-33725 AF488) diluted in 1% BSA. After washing, Alexa Fluor 488-conjugated secondary antibody (1:500) was applied for 2 hours at room temperature, and nuclei were counterstained with DAPI. Flatmounts were mounted using ProLong Gold Antifade mounting medium and imaged with a Cytation 7 microscope. ZO-1 staining intensity and tight junction morphology were quantified using ImageJ.

### Transmission Electron Microscopy

Eyes were enucleated immediately following euthanasia and processed for transmission electron microscopy as previously described with minor modifications. Briefly, eyes were punctured at the corneal limbus and immersion-fixed in **2.5% glutaraldehyde in 0.1 M cacodylate buffer (pH 7.4)** at 4 °C overnight. Following fixation, posterior eyecups were dissected to isolate the retinal tissue and rinsed thoroughly in cacodylate buffer. Samples were post-fixed in **1% osmium tetroxide** for 1 h at room temperature, rinsed, and en bloc stained with **2% aqueous uranyl acetate**. Tissues were then dehydrated through a graded ethanol series and embedded in epoxy resin. Ultrathin sections were cut using an ultramicrotome and collected on copper grids. Sections were counterstained with **uranyl acetate and lead citrate** prior to imaging. Ultrastructural imaging was performed using a transmission electron microscope operated at **80–120 kV**. Images were acquired to assess the ultrastructure of the **retinal pigment epithelium (RPE), photoreceptor inner and outer segments**, and the **choroidal vasculature**, with particular attention to mitochondrial morphology and sub-RPE basal laminar deposits. Representative images were captured at multiple magnifications for each retinal layer.

### Quantification of L-Serine

D- and L-Serine levels were quantified using the DL-Serine Assay Kit (Sigma-Aldrich, Cat# MAK352), following the manufacturer’s protocol in the serum of mice and ARPE-19 cells treated with/without L-serine. Briefly, RPE sheets were carefully dissected and homogenized in Serine Assay Buffer using a homogenizer on ice. Cultured human ARPE-19 and iPSC-RPE cells were lysed in cold Serine Assay Buffer and processed similarly. All samples were deproteinized with trichloroacetic acid, incubated at 37°C for 15 minutes, and then centrifuged through 5000 MWCo Corning Spin-X® UF concentrators (10,000 × g, 10 min). Lysates were incubated with the kit’s enzyme mix for 1 hour. Fluorescence was recorded at λ_ex/em 535/587 nm.

### Reactive oxygen species measurement

Dihydroethidium (DHE) fluorescence-based ROS detection kit was used to determine ROS generated. 40,000 ARPE-19 cells were plated on dark walled 96 well plates and treated with siPINK1 or siScramble and subject to serine supplementation as mentioned above. Cells were then incubated with 5uM DHE for 30 minutes at 37C protected from light. Fluorescence was measured using a Cytation 7 plate reader with λex/em 480/570 nm.

### Mitochondrial superoxide measurement

ARPE-19 cells were plated on dark walled 96 well plates and treated with scramble siRNA or Pink1 siRNA with and without serine for 5 days. Mitochondrial superoxide levels were measured using MitoSOX Red (Invitrogen M36008). Stock solution of 5 mM was made in DMSO. They were then washed with PBS and treated with a working solution of 500 nM for 1 hour in HBSS with calcium and magnesium. Cells were then washed with HBSS. Cells were then treated with 1ug/mL Hoechst 33342 for 5 minutes and fluorescence was imaged with Cytation 7 microscope at λex/em 396/610 nm. Images were then analyzed using ImageJ.

### Mitochondrial membrane potential measurement

ARPE-19 cells were plated on dark walled 96 well plates and treated with scramble siRNA or Pink1 siRNA with and without serine for 5 days. Mitochondrial membrane potential was measured using tetramethylrhodamine ethyl ester (Invitrogen T669). Stock solution of 1 mM was made in DMSO. They were then washed with PBS and treated with a working solution of 1uM for 1 hour in HBSS with calcium and magnesium. Cells were then washed and treated with 1ug/mL Hoechst 33342 for 5 minutes and fluorescence was imaged with Cytation 7 microscope at λ_ex = 549 nm and λ_em = 574 nm. Images were then analyzed using ImageJ.

### Seahorse

Mitostress assay recordings of oxygen consumption rate (OCR) and (ECAR) were performed on a Seahorse XF Pro analyzer. Cells were treated with either siScr or siPink1 and 40,000 cells were seeded onto the 96 well microplate. They were then treated with 200uM serine for 5 days. On day 5, cells were washed with XF DMEM media supplemented with 5mM glucose, 1mM pyruvate and 2mM glutamine. Three compounds (oligomycin, 2uM; FCCP, 2uM; Antimycin A/Rotenone, 10uM) which alter mitochondrial activity were prepared in the same XF DMEM media. These compounds were then added onto the cartridge and loaded into the Seahorse XF Pro. For each measurement, 6 total readings were recorded at an interval of 3 minutes each with a total of 18 readings for each replicate.

### Mice

Mice were bred at the Emory Eye Center according to the animal welfare guidelines outlined in the Emory University Institutional Animal Care and Use Committee (IACUC). All procedures were approved by the IACUC prior to experimentation. Mice were housed in 12 hour dark-light cycles and were maintained on a commercially available chow diet. Wildtype C57BL/6 and Pink1^-/-^ were bred in our colony. For *in vivo* experiments, mice were aged for 12 months followed by 6 months of serine treatment (o.5%w/v) in ad libitum drinking water. The mice were then screened for rd10 and rd8 retinal degeneration mutations. Both male and female mice were used to account for possible sex-based differences.

### Serine in drinking water

L-serine (BeanTown Chemical, NH, Cat# 128350) was dissolved in autoclaved drinking water at 0.5% w/v. Drinking water was provided *ad libitum* for a duration of 6 months. Serine water was changed every week.

### Fundus Imaging

Mice were anesthetized with 4mg/mL ketamine and 1mg/mL xylazine. Eyes were then covered with gel and imaged on Micron IV imaging system.

### H&E stain

Eyes were freeze substituted with 97% methanol and 3% glacial acetic acid for 1 week. They were dehydrated in gradient ethanols and xylenes. Eyes were then embedded in paraffin and 5um sections were rehydrated and stained with Mayer’s Hematoxylin for 1 minute. Slides were washed in tap water and then stained with eosin for 1 minute, dehydrated in gradient ethanols and mounted.

## GRAPHICAL ABSTRACT

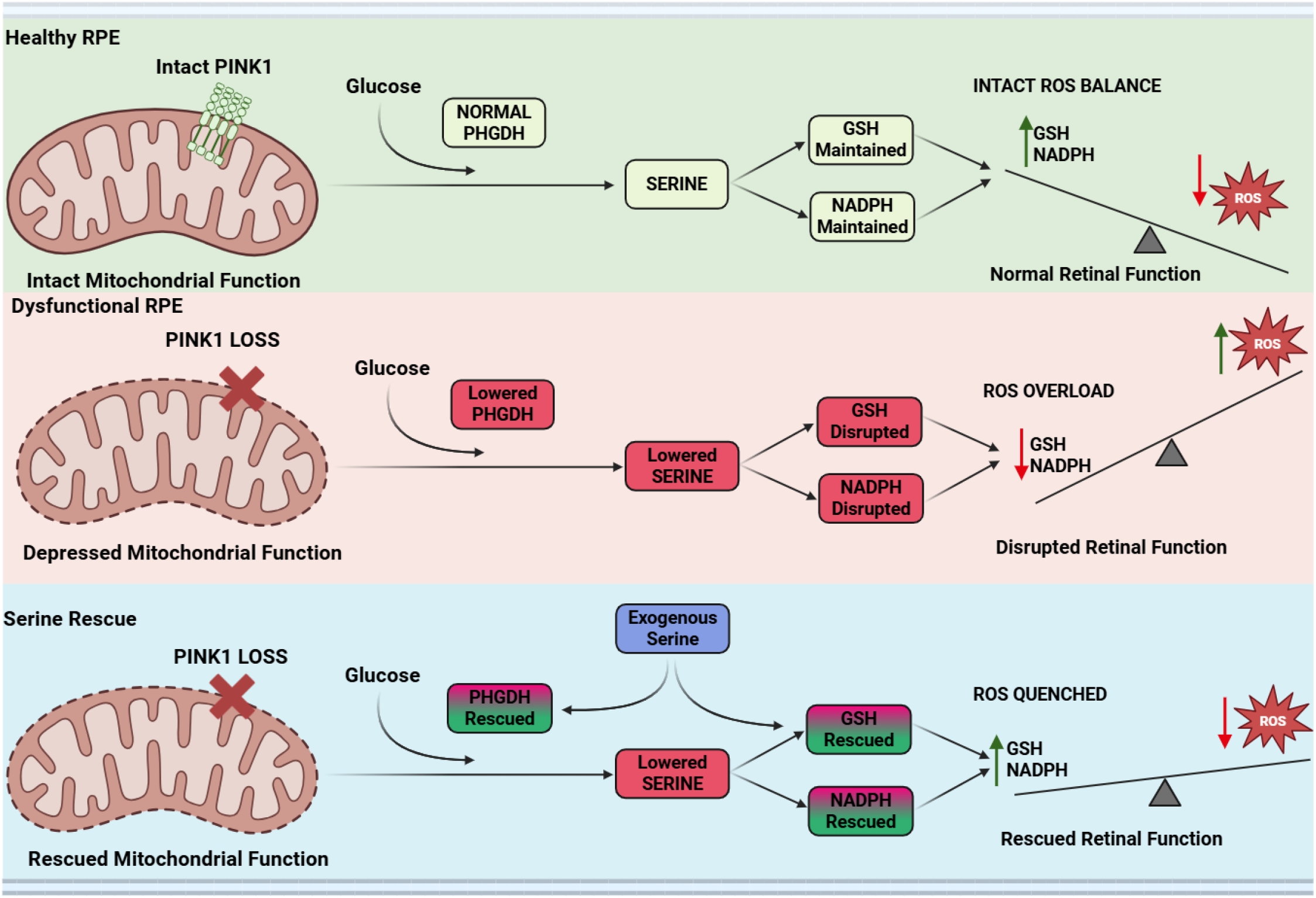

## REFERENCES

1. Brown GC, Brown MM, Sharma S, Stein JD, Roth Z, Campanella J, Beauchamp GR. The burden of age-related macular degeneration: a value-based medicine analysis. Trans Am Ophthalmol Soc. 2005;103:173–184 184–176.

2. Zajac-Pytrus HM, Pilecka A, Turno-Krecicka A, Adamiec-Mroczek J, Misiuk-Hojlo M. The Dry Form of Age-Related Macular Degeneration (AMD): The Current Concepts of Pathogenesis and Prospects for Treatment. Adv Clin Exp Med. 2015;24:1099–1104.

3. Datta S, Cano M, Satyanarayana G, Liu T, Wang L, Wang J, Cheng J, Itoh K, Sharma A, Bhutto I, et al. Mitophagy initiates retrograde mitochondrial-nuclear signaling to guide retinal pigment cell heterogeneity. Autophagy. 2023. doi: 10.1080/15548627.2022.2109286.

4. Datta S, Cano M, Ebrahimi K, Wang L, Handa JT. The impact of oxidative stress and inflammation on RPE degeneration in non-neovascular AMD. Prog Retin Eye Res. 2017;60:201–218.

5. Terluk MR, Kapphahn RJ, Soukup LM, Gong H, Gallardo C, Montezuma SR, Ferrington DA. Investigating mitochondria as a target for treating age-related macular degeneration. J Neurosci. 2015;35:7304–7311. doi: 10.1523/JNEUROSCI.0190-15.2015

6. Wang S, Long H, Hou L, Feng B, Ma Z, Wu Y, Zeng Y, Cai J, Zhang DW, Zhao G. The mitophagy pathway and its implications in human diseases. Signal Transduct Target Ther. 2023;8:304. doi: 10.1038/s41392-023-01503-7

7. Lu Y, Li Z, Zhang S, Zhang T, Liu Y, Zhang L. Cellular mitophagy: Mechanism, roles in diseases and small molecule pharmacological regulation. Theranostics. 2023;13:736–766. doi: 10.7150/thno.79876

8. Karunadharma PP, Nordgaard CL, Olsen TW, Ferrington DA. Mitochondrial DNA damage as a potential mechanism for age-related macular degeneration. Invest Ophthalmol Vis Sci. 2010;51:5470–5479.

9. Kaarniranta K, Uusitalo H, Blasiak J, Felszeghy S, Kannan R, Kauppinen A, Salminen A, Sinha D, Ferrington D. Mechanisms of mitochondrial dysfunction and their impact on age-related macular degeneration. Prog Retin Eye Res. 2020;79:100858.

10. Yang M, Vousden KH. Serine and one-carbon metabolism in cancer. Nat Rev Cancer. 2016;16:650–662.

11. Reid MA, Allen AE, Liu S, Liberti MV, Liu P, Liu X, Dai Z, Gao X, Wang Q, Liu Y, et al. Serine synthesis through PHGDH coordinates nucleotide levels by maintaining central carbon metabolism. Nat Commun. 2018;9:5442.

12. Zhou X, He L, Wu C, Zhang Y, Wu X, Yin Y. Serine alleviates oxidative stress via supporting glutathione synthesis and methionine cycle in mice. Mol Nutr Food Res. 2017;61.

13. Wang F, Zhou H, Deng L, Wang L, Chen J, Zhou X. Serine Deficiency Exacerbates Inflammation and Oxidative Stress via Microbiota-Gut-Brain Axis in D-Galactose-Induced Aging Mice. Mediators Inflamm. 2020:5821428.

14. Sinha T, Ikelle L, Naash MI, Al-Ubaidi MR. The Intersection of Serine Metabolism and Cellular Dysfunction in Retinal Degeneration. Cells. 2020. doi: 10.3390/cells9030674.

15. Zhou X, He L, Zuo S, Zhang Y, Wan D, Long C, Huang P, Wu X, Wu C, Liu G, Yin Y. Serine prevented high-fat diet-induced oxidative stress by activating AMPK and epigenetically modulating the expression of glutathione synthesis-related genes. Biochim Biophys Acta Mol Basis Dis. 2018;1864:488–498. doi: 10.1016/j.bbadis.2017.11.009

16. Gantner ML, Eade K, Wallace M, Handzlik MK, Fallon R, Trombley J, Bonelli R, Giles S, Harkins-Perry S, Heeren TFC, et al. Serine and Lipid Metabolism in Macular Disease and Peripheral Neuropathy. N Engl J Med. 2019;381:1422–1433. doi: 10.1056/NEJMoa1815111

17. Rodriguez AE, Ducker GS, Billingham LK, Martinez CA, Mainolfi N, Suri V, Friedman A, Manfredi MG, Weinberg SE, Rabinowitz JD, Chandel NS. Serine Metabolism Supports Macrophage IL-1β Production. Cell Metab. 2019. doi: 10.1016/j.cmet.2019.01.014.

18. Tajan M, Hennequart M, Cheung EC, Zani F, Hock AK, Legrave N, Maddocks ODK, Ridgway RA, Athineos D, Suarez-Bonnet A, et al. Serine synthesis pathway inhibition cooperates with dietary serine and glycine limitation for cancer therapy. Nat Commun. 2021;12:366.

19. Eade K, Gantner ML, Hostyk JA, Nagasaki T, Giles S, Fallon R, Harkins-Perry S, Baldini M, Lim EW, Scheppke L, et al. Serine biosynthesis defect due to haploinsufficiency of PHGDH causes retinal disease. Nat Metab. 2021;3:366–377. doi: 10.1038/s42255-021-00361-3

20. Lim EW, Fallon RJ, Bates C, Ideguchi Y, Nagasaki T, Handzlik MK, Joulia E, Bonelli R, Green CR, Ansell BRE, et al. Serine and glycine physiology reversibly modulate retinal and peripheral nerve function. Cell Metab. 2024;36:2315–2328 e2316. doi: 10.1016/j.cmet.2024.07.021

21. Lowy Medical Research Institute. (2021). Phase 2a study of the effect of serine supplementation and fenofibrate treatment on serum deoxysphinganine levels in patients with macular telangiectasia (MacTel) type 2 (SAFE study) (ClinicalTrials.gov Identifier NCT04907084).

22. Massachusetts General Hospital. (2012). Randomized trial of L-serine in patients with hereditary sensory and autonomic neuropathy type 1 (HSAN1) (ClinicalTrials.gov Identifier NCT01733407).

23. University College London. (2023). Hereditary sensory neuropathy serine trial (SENSE trial) (ClinicalTrials.gov Identifier NCT06113055).

