## Supplementary figures and images for "Dietary serine protects the retinal pigmented epithelium by blunting reactive oxygen species in dry age-related macular degeneration"

### Supplementary Material

#### PINK1 Manuscript

##### Supplementary Figures

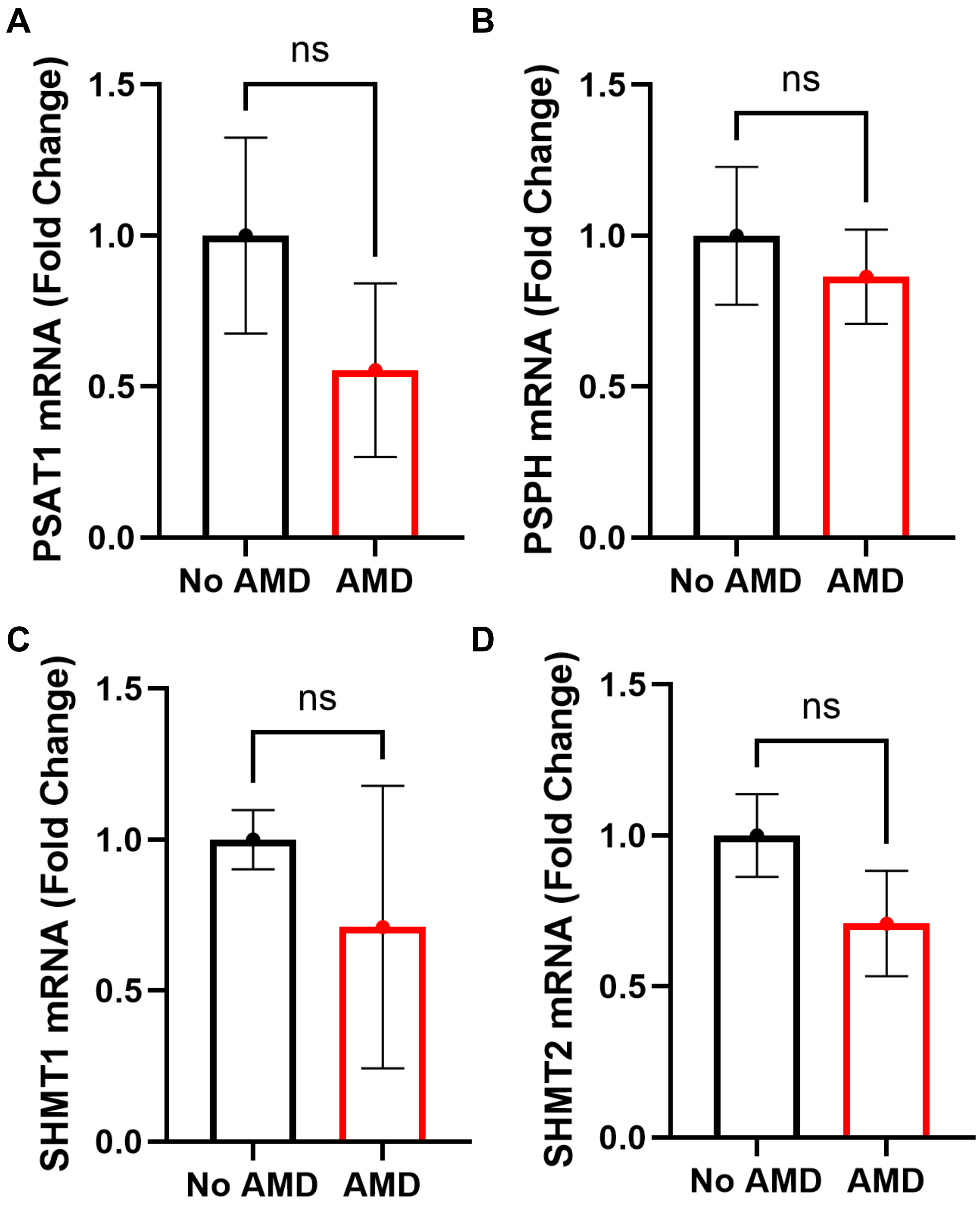

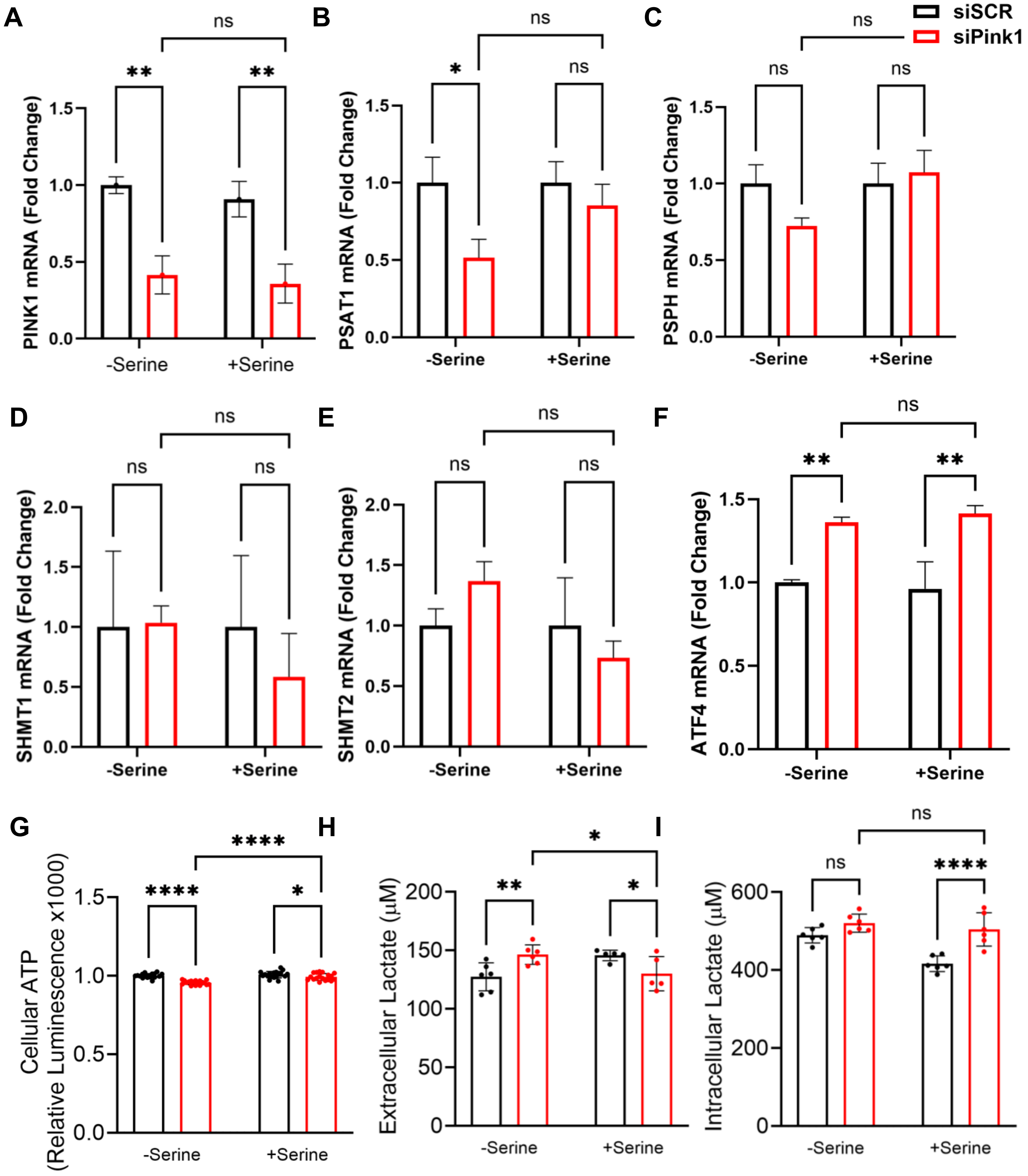

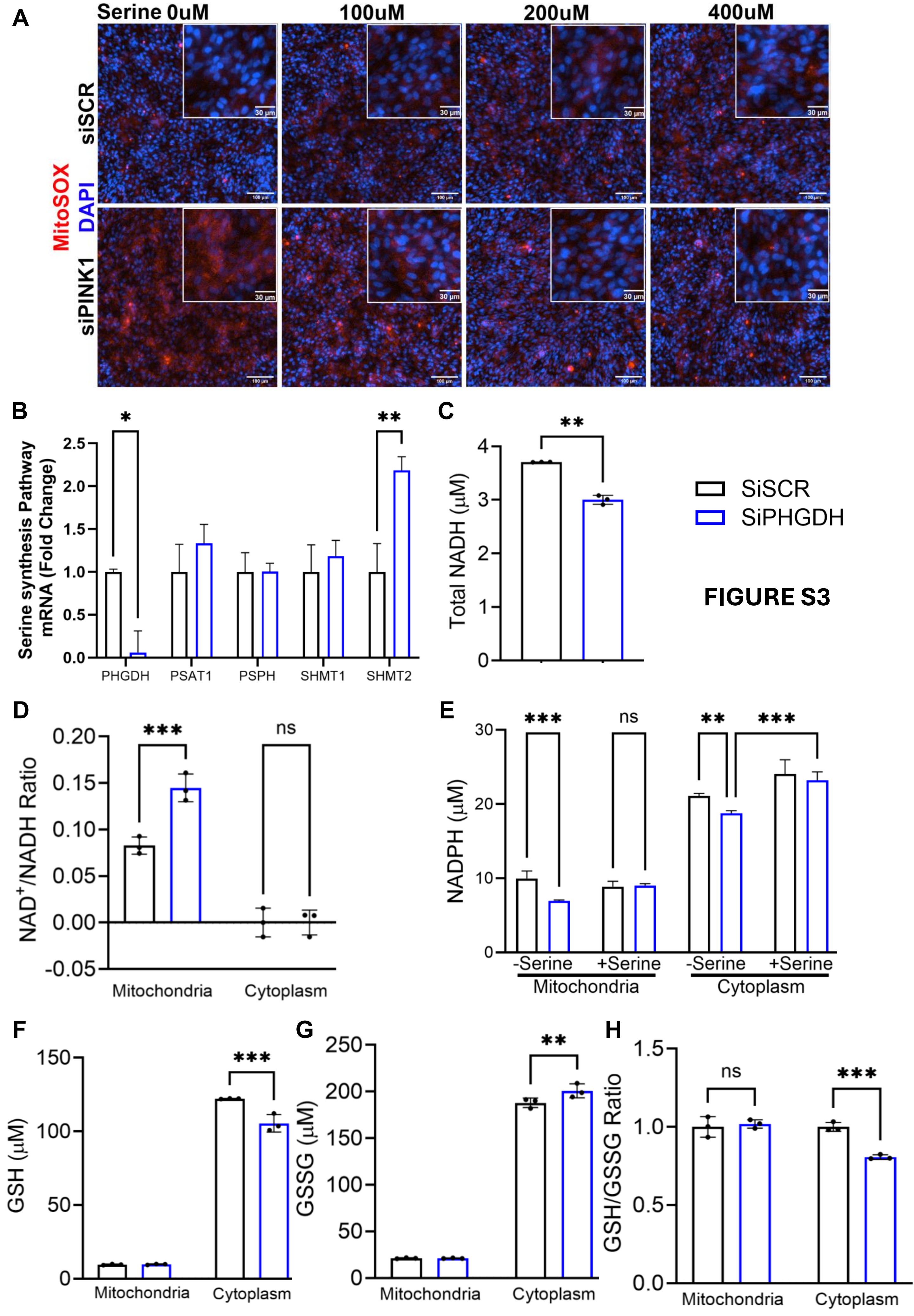

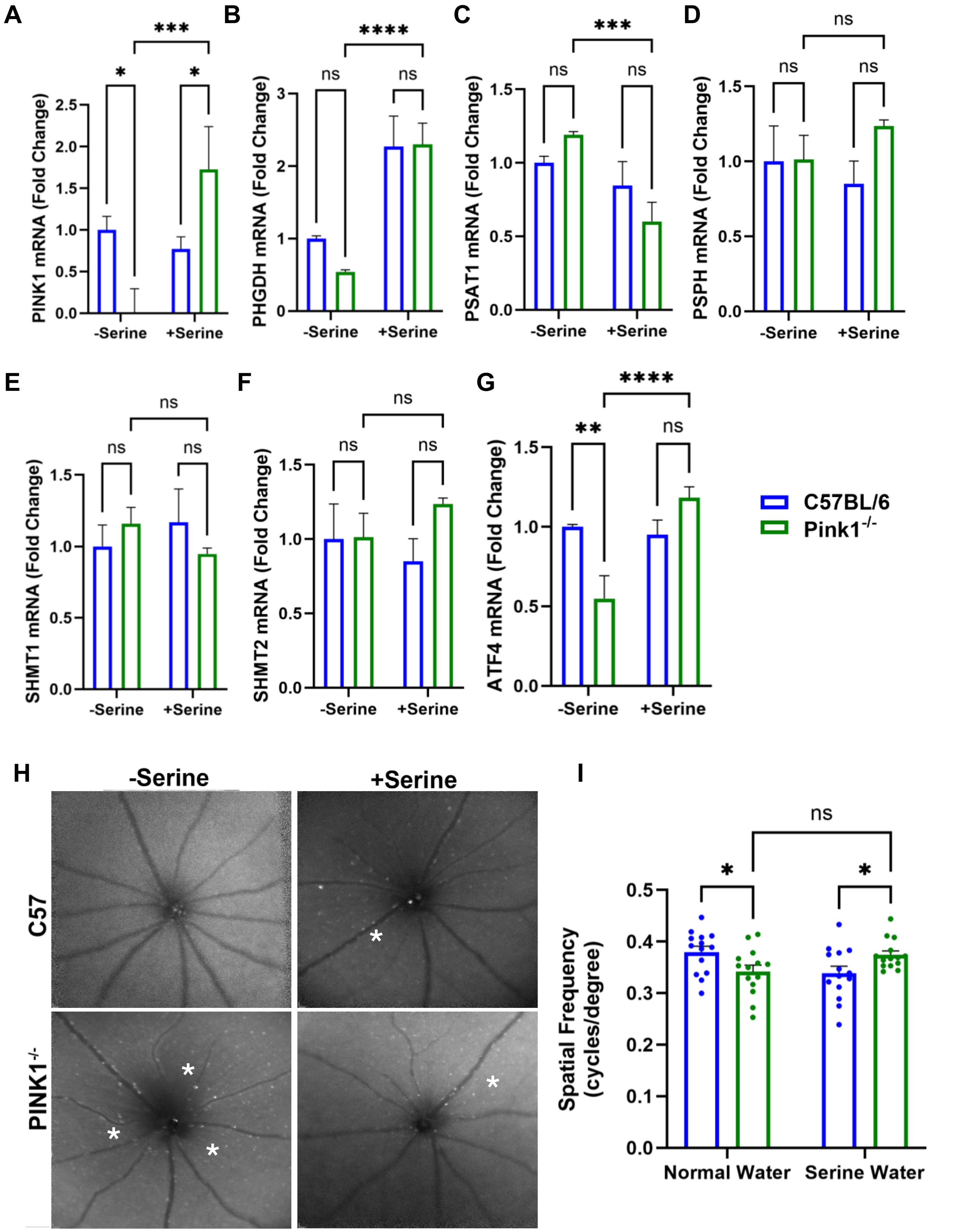
