## Supplementary Figure Legends for "Dietary serine protects the retinal pigmented epithelium by blunting reactive oxygen species in dry age-related macular degeneration"

**Figure S1. Serine synthesis pathway genes are unaltered AMD patient derived iPSC-RPE**
RNA was isolated and qPCR was performed on iPSCs from AMD patients and non-AMD controls for the following serine synthesis pathway genes (**A**) Phosphoserine aminotransferase (PSAT1) (**B**) Phosphoserine phosphatase (PSPH) (**C**) Cytosolic serine hydroxymethyl transferase 1 (SHMT1) (**D**) Mitochondrial serine hydroxymethyl transferase 2 (SHMT2).

**Figure S2. Pink1 loss affects serine synthesis pathway genes and glycolytic end products**
ARPE-19 cells treated with siPink1 and supplemented with serine for 5 days. (**A-E**) qPCR was performed for Pink1, PSAT1, PSPH, SHMT1, SHMT2 mRNA levels. (**F**) Total cellular ATP levels were assessed using Cell-titer-Glo (G9241). (**G**) Secreted and (**H**) intracellular lactate levels were measured in these cells with Lactate-Glo (J5021). (**I**) Total pyruvate levels were quantified fluorometrically using a kit (Sigma MAK332). (Data are presented as mean +/- SEM from n = 6 replicates per group. Statistical significance was determined using two-way ANOVA with siPINK1 (knockdown vs. control) and +/- serine supplementation, followed by Tukey's post hoc multiple comparisons test. (*p<0.05; **p<0.005; ****p<0.0001)

**Figure S3. Serine ameliorates mitochondrial superoxides and rescues ROS management systems**
(**A**) ARPE-19 cells were treated with 20pmol siPINK1 for 5 days followed by serine supplementation at 100uM, 200uM and 400uM. MitoSOX stain was performed to assess mitochondrial superoxide accumulation. (**B**) qPCR analyses serine synthesis pathway genes. (**C**) Total NADH was measured fluorometrically using a NAD+/NADH kit (Abcam 176723) (**D**) Mitochondrial and cytoplasmic fractions were isolated and NAD+/NADH ratios were measured in both fractions (**E**) NADPH levels in these fractions were assessed with serine supplementation. (**F**) Total reduced glutathione (GSH) and (**G**) Total oxidized glutathione (GSSG) (**H**) GSH/GSSG ratio were measured in the cytoplasmic and mitochondrial fractions using a kit (MedChemExpress HY-K0311). Data are presented as mean +/- SEM from n = 3 replicates per group. Statistical significance was determined using two-way ANOVA with siPHGDH (knockdown vs. control) and +/- serine supplementation, followed by Tukey's post hoc multiple comparisons test. (*p<0.05; **p<0.005; ***p<0.0005)

**Figure S4.**
RNA was isolated from RPE explants from C57BL/6 and Pink1-/- mice treated with either normal water or serine water qPCR was performed for (**A**) PINK1 (**B**) PHGDH (**C**) PSAT1 (**D**) PSPH (**E**) SHMT1 (**F**) SHMT (**G**) ATF4 (**H**) IR field fundus photography was performed. White stars represent hyper-reflective spots in the fundus. (**I**) Optokinetic responses were measured and spatial frequencies were recorded as a measure of contrast sensitivity. Data are presented as mean +/- SEM from n = 6 replicates per group. Statistical significance was determined using two-way ANOVA with C57BL/6 vs Pink1-/- and +/- serine supplementation, followed by Tukey's post hoc multiple comparisons test. (*p<0.05; ***p<0.0005; ****p<0.0001)
